# Dynamic activity of *Erg* promotes aging of the hematopoietic system

**DOI:** 10.1101/2025.01.23.634563

**Authors:** Mayuri Tanaka-Yano, Wade W. Sugden, Dahai Wang, Brianna Badalamenti, Parker Côté, Diana Chin, Julia Goldstein, Stephan George, Gabriela F. Rodrigues-Luiz, Edroaldo Lummertz da Rocha, Hojun Li, Trista E. North, Berkley E. Gryder, R. Grant Rowe

**Affiliations:** Stem Cell Program and Stem Cell Transplantation Programs, Boston Children’s Hospital, Boston, MA; Department of Hematology/Oncology, Boston Children’s Hospital, Boston, MA; Koch Institute for Integrative Cancer Research, Massachusetts Institute of Technology, Cambridge, MA; Department of Genetics & Genome Sciences, Case Western Reserve University, Cleveland, OH; Federal University of Santa Catarina, Florianópolis, Brazil; Division of Hematology/Oncology, Department of Pediatrics, University of California, San Diego, La Jolla, CA; Harvard Medical School, Boston, MA; Department of Pediatric Oncology, Dana-Farber Cancer Institute, Boston, MA

## Abstract

Hematopoiesis changes to adapt to the physiology of development and aging. Temporal changes in hematopoiesis parallel age-dependent incidences of blood diseases. Several heterochronic regulators of hematopoiesis have been identified, but how the master transcription factor (TF) circuitry of definitive hematopoietic stem cells (HSCs) adapts over the lifespan is unknown. Here, we show that expression of the ETS family TF *Erg* is adult-biased, and that programmed upregulation of *Erg* expression during juvenile to adult aging is evolutionarily conserved and required for complete implementation of adult patterns of HSC self-renewal and myeloid, erythroid, and lymphoid differentiation. *Erg* deficiency maintains fetal transcriptional and epigenetic programs, and persistent juvenile phenotypes in *Erg* haploinsufficient mice are dependent on deregulation of the fetal-biased TF *Hmga2*. Finally, *Erg* haploinsufficiency in the adult results in fetal-like resistance to leukemogenesis. Overall, we identify a mechanism whereby HSC TF networks are rewired to specify stage-specific hematopoiesis, a finding directly relevant to age-biased blood diseases.

**SUMMARY:** The hematopoietic system undergoes a process of coordinated aging from the juvenile to adult states. Here, we find that expression of ETS family transcription factor Erg is temporally regulated. Impaired upregulation of *Erg* during the hematopoietic maturation results in persistence of juvenile phenotypes.

## INTRODUCTION

The definitive hematopoietic system undergoes a stereotypic process of maturation and aging over time from the fetal to adult state characterized by alterations in mature cell output on schedule with age-related changes in overall physiology^1^. These temporal changes in hematopoiesis likely underpin age biases in the incidence of many blood diseases. At the cellular level, changes in mature cell production are governed by variation in self-renewal, lineage biases, transcriptional priming, and fate decisions at the level of HSCs^2–7^. At the molecular level, these age-related changes in stem cell properties are regulated by highly conserved heterochronic factors that coordinate the timing of morphogenetic events to maintain programmed schedules of development, maturation, and aging^5,8–10^. The RNA binding protein Lin28b is a master heterochronic regulator within the hematopoietic system^11^. Lin28b expression is fetal-biased, and its developmental downregulation enables the processing and maturation of mRNAs and microRNAs that implement varied aspects of adult hematopoiesis^5,8–10,12–16^.

TFs play central roles in hematopoiesis by defining stem and effector cell states, promoting or restraining self-renewal, and regulating fate choices; their dysregulation leads to dysfunction of HSCs in disease states such as bone marrow failure disorders, myelodysplastic syndrome, and leukemia^17–19^. Given their central roles in hematopoiesis, several TFs have been implicated in regulation of the age state of the definitive blood forming system, but how key heterochronic factors impact the core HSC TF circuitry remains unclear^2,20,21^. In our previous work, we found that Lin28b regulates the expression of master hematopoietic TFs via the Polycomb repressive complex 1 (PRC1) component Cbx2, providing a mechanism by which Lin28b exerts global effects on hematopoiesis via regulation of TF expression^10^.

Several erythroblast transformation-specific (ETS) family TFs play central roles in hematopoiesis. *Erg* is an ETS family TF that is essential for specification of HSCs during development, and perturbation of Erg function has been implicated in several forms of leukemia in children and adults^22–26^. We previously found that *Erg* expression is repressed by PRC1/Cbx2 downstream of Lin28b in fetal HSCs and progenitors as evidence for potential heterochronic functions of *Erg* in blood formation^10^. In further support of this idea, although complete loss of *Erg* results in failure of definitive hematopoiesis, deletion of one allele of *Erg* in mice is tolerated during development but results in perturbed of HSC maintenance and self-renewal in the adult^22,27–30^. Using conditional knockout models, Erg has otherwise been implicated in murine B-cell specification^31^.

Here, we find that expression of *Erg* in definitive hematopoietic stem and progenitor cells (HSPCs) is temporally regulated, increasing during aging from the juvenile to adult state. While *Erg* heterozygosity has minimal impact on fetal hematopoiesis, diminished *Erg* dosage results in inappropriate persistence of fetal hematopoietic phenotypes into adulthood at the levels of HSPCs, lineage-restricted progenitors, and mature effector cells. Erg regulates transcriptional and epigenetic programs controlling the juvenile to adult transition in the hematopoietic system, with inadequate downregulation of the fetal factor *Hmga2* in adult *Erg* haploinsufficient mice implicated in persistent fetal phenotypes. Blunting of *Erg* expression in adult HSPCs causes resistance of leukemic transformation, positing age-related upregulation of *Erg* as contributing to the increased incidence of myeloid malignancies in older adults. Together, our results identify a role of Erg in heterochronic control of hematopoietic maturation, a finding highly relevant to Erg-associated blood diseases and broaden our understanding of paradigms of hematopoietic regulation by TFs by incorporating the dimension of real time remodeling of HSPC TF networks.

## RESULTS

### Dynamic expression and activity of ETS factors during the juvenile to adult hematopoietic transition

To understand regulation of juvenile to adult hematopoietic aging, we analyzed two independent datasets comparing chromatin accessibility profiles (assay of transposase accessible chromatin with sequencing (ATAC-seq)) between definitive multipotent midgestation murine fetal and adult HSPCs^14,32^. To uncover candidate TFs promoting hematopoietic aging, we defined TF motifs enriched in chromatin regions specifically accessible in adult HSPCs. We found that ETS family motifs were the most significantly adult-biased in both datasets (Figure 1A). In a previous study, we observed developmentally regulated expression of the ETS factor *Erg* downstream of the heterochronic Lin28b-*let-7*-Cbx2 axis^10^. Here, we confirmed that expression of *Erg* increased in murine Lineage^-^ Sca-1^+^ ckit^+^ (LSK) multipotent HSPCs during the fetal (embryonic age 14.5 fetal liver (E14.5 FL) to adult (∼12 weeks) transition (Figure 1B). Similarly, in a published dataset, we found that expression of *ERG* increases during the fetal to adult transition in human HSPCs (Figure 1C)^21^. Furthermore, using a dataset from zebrafish hematopoietic tissues, we observed that *erg* expression was increased in adult kidney marrow (KM; bone marrow ortholog) relative to caudal hematopoietic tissue (CHT; FL ortholog; Figure 1D)^33^. Finally, using a single cell transcriptomic dataset, we found that *ERG* expression increased in HSPCs over the human lifetime, and markedly into advanced age (Figure 1E)^34^. Consistent with these findings, the murine *Erg* locus showed a higher degree of chromatin accessibility in adult HSCs compared to fetal (Figure 1F)^14^. Together, these findings show that *Erg* undergoes a conserved increased in expression with aging and suggest that its activity increases during the juvenile to adult hematopoietic transition.

**Figure 1.**
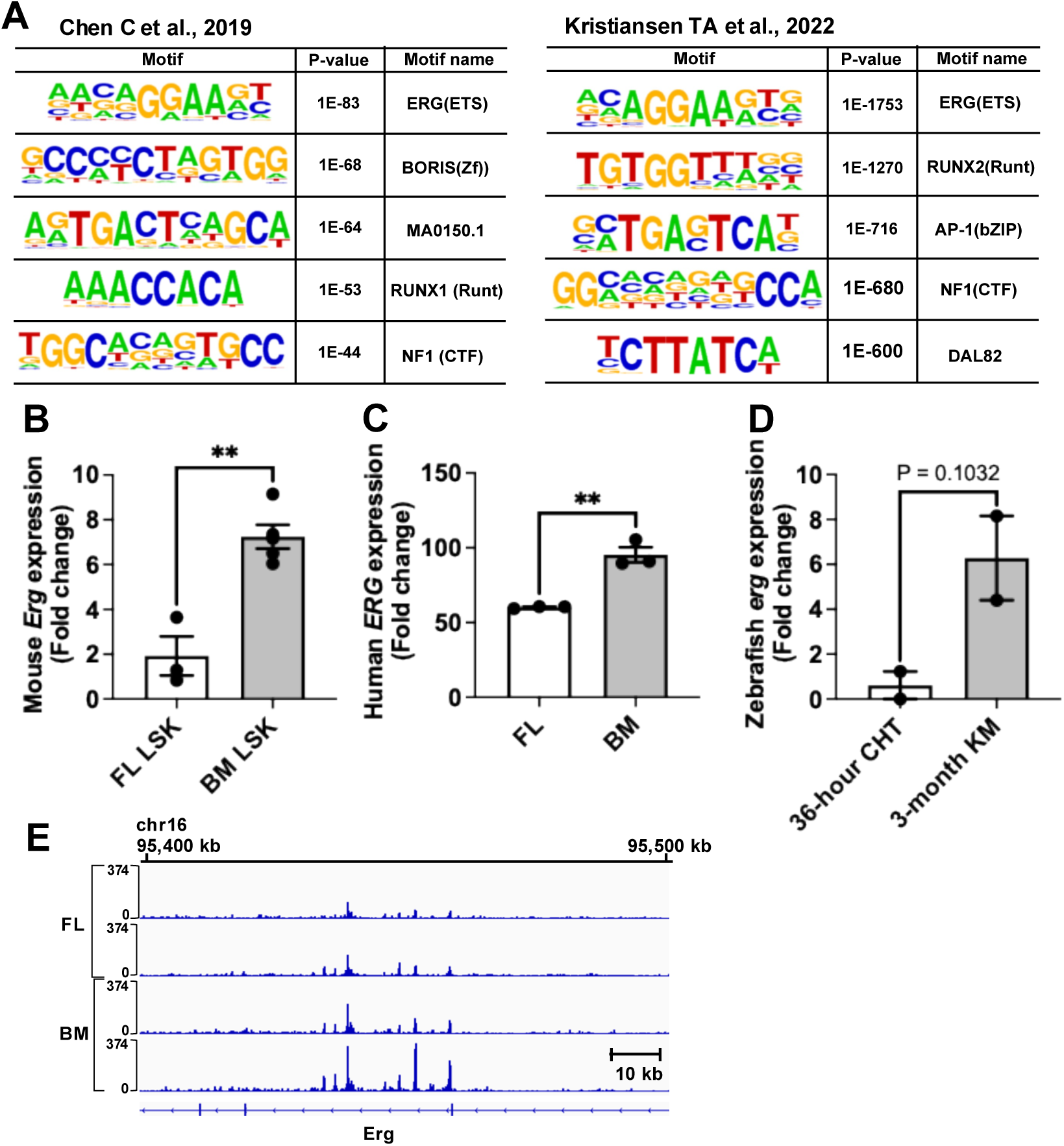
Adult biased *Erg* expression and ETS factor motif usage. (A) HOMER analysis of motifs enriched in adult HSPC-specific accessible chromatin regions (relative to fetal) in the indicated datasets^14,32,76^. (B) Mouse *Erg* expression level in LSK (Lin^-^Sca1^+^cKit^+^) from FL (E14.5) and BM (12-week-old) by qPCR (n = 3-5 per group). Data are shown as mean ± SEM. (C) Human *ERG* expression in FL and BM HSPCs by RNA-seq (n = 3). Data are shown as median ± SEM^21^. (D) Zebrafish *erg* expression level in 36-hr caudal hematopoietic tissue (CHT) and 3-month KM by RNA-seq (n = 2). Data are shown as mean ± SEM^33^. (E) Single cell-level expression of *ERG* at the indicated age stages, with heatmap showing - log_10_ transformed p-values for the indicated comparisons (Wilcoxon rank-sum test)^34^. (F) Visualization of ATAC-seq tracks at the *Erg* locus in fetal and adult HSPCs^14^. In all panels, comparisons by Student’s t test; \*\**P* < 0.01.

### Full dosing of Erg is required for juvenile to adult hematopoietic aging

The observed adult-skewed *Erg* expression as well as previously reported hematopoietic phenotypes in *Erg* haploinsufficient adult mice led us to hypothesize that the temporally programed increase in *Erg* expression is important for full implementation of the adult HSPC state. To test this, we generated Vav-Cre; *Erg*^flox/+^ mice (*Erg*^+/-^) to blunt the temporal increase in *Erg* expression that occurs during the juvenile to adult HSPC transition^35^. This resulted in an approximately 40% reduction in *Erg* expression in mature adulthood (Figure 2A).

**Figure 2.**
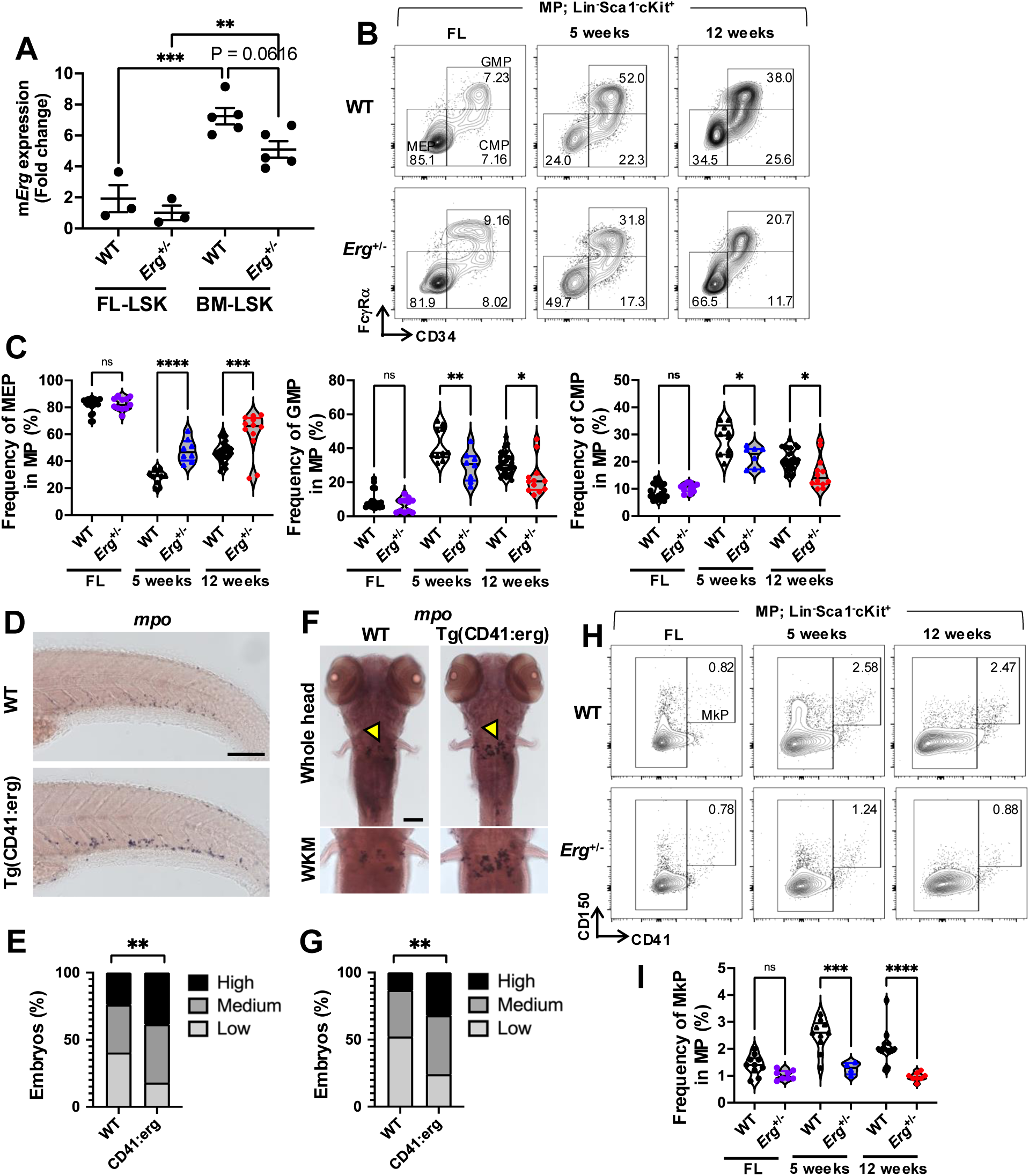
*Erg* regulates maturation of the myeloerythroid compartment. (A) qPCR analysis of *Erg* expression in FL and BM from WT and *Erg*^+/-^ mice (n = 3-5 biological replicates per group). The results were pooled from three independent experiments. (B) Representative MEP, GMP, and CMP distributions among MPs in FL, 5-, and 12-week-old BM. Populations were gated on Lin^-^Sca1^-^cKit^+^. (C) Frequency of MEP, GMP, and CMP in MP from FL, 5-, and 12-week-old BM (n = 8-28 biological replicates per group). The results were pooled from six independent experiments. (D) Whole-mount in situ hybridization (WISH) for mpo at 3 days post fertilization (dpf) in caudal hematopoietic tissue (CHT) in WT and Tg(CD41:*erg*) zebrafish. Scale bar, 100 µm. (E) Qualitative phenotypic distribution plot for high, medium and low *mpo* expression in embryos scored in D (WT, n = 88, Tg, n = 83). The results were pooled from three independent experiments. (F) WISH for *mpo* at 5 dpf in WT and Tg(CD41:*erg*) zebrafish. Yellow arrowhead indicates cells stained in the whole KM (WKM), shown in inset. Scale bar, 100 µm. (G) Phenotypic distribution plot for embryos scored in F (WT, n = 46, Tg, n = 50). The results were pooled from two independent experiments. (H) Representative distributions of MkP in FL, 5-, and 12-week-old BM. (I) Frequency of MkP from FL, 5-, and 12-week-old BM (n = 4-12 per group). The results were pooled from six independent experiments. Data are shown as mean ± SEM. Statistical tests were one-way ANOVA with Tukey’s multiple comparisons test (A, C, I) or Chi-square test (E, G). \**P* < 0.05; \*\**P* < 0.01; \*\*\**P* < 0.001; \*\*\*\**P* < 0.0001.

We then examined hallmarks of the juvenile to adult definitive hematopoietic transition (Figure S1A). Hematopoietic aging is associated with a skew toward myeloid output and away from juvenile erythroid-dominated output which manifests in an increased proportion of granulocyte monocyte progenitors (GMPs) relative to megakaryocyte erythroid progenitors (MEPs; Figure S2A)^9^. While we did not observe disruption of the balance of these myeloid progenitors (MPs) in the FL, in the young (5-week) and mature (12-week) adult bone marrow (BM) hematopoietic compartment, *Erg*^+/-^ mice showed blunting of the programmed increase in GMPs and decrease in MEPs compared to littermates with an overall juvenile-like MP distribution (Figure 2B-C). We observed an analogous effect in zebrafish models, where overexpression of *erg* under the HSPC/thrombocyte-restricted promoter of *cd41* resulted in myeloid skewing in the CHT at 3 days post fertilization (dpf) with expansion of *mpo*-expressing cells, which persisted in the KM stage at 5 dpf (Figure 2D-G). Furthermore, in the myeloerythroid lineage, the aging results in emergence of Lineage^-^Sca1^-^ckit^+^CD150^+^CD41^+^ megakaryocyte progenitors (MkPs; Figure S2B)^14^. We found that MkP emergence is impaired in *Erg*^+/-^ adults, which is associated with alterations in megakaryocyte (MK) subpopulations postnatally, specifically, blunted temporal increases in MK stages II and V (Figure 2H-I, S1C-H).

Mature lymphoid populations have varied developmental origins. B1 and marginal zone (MZ) B-cells have origins in prenatal hematopoiesis, while B2 and follicular (Fo) B-cells arise postnatally (Figure S3A)^5^. We found that *Erg*^+/-^ adults showed an increase in MZ and diminishment of Fo B-cells compared to littermate controls, although the B1/B2 balance was not significantly altered at steady state (Figure 3A-D). However, ex vivo culture of *Erg*^+/-^ and littermate multipotent LSK HSPCs under lymphopoietic conditions revealed skewing toward fetal-like B-cell output in *Erg*^+/-^ LSKs, as did transplantation of LT-HSCs (Figure 3E-F, S3B). In the zebrafish, we observed a role for *erg* in regulating the juvenile lymphoid bias, with ectopic *ERG* expression skewing away from lymphoid output and, conversely, *erg* morpholino knockdown promoting lymphopoiesis (Figure 3G-H). Together, these results demonstrate that temporal upregulation of *Erg* is required for full implementation of myeloid-biased adult hematopoiesis relative to lymphoid- and erythroid-biased juvenile hematopoiesis, and that *Erg* plays a role in adult B-cell specification^36^.

**Figure 3.**
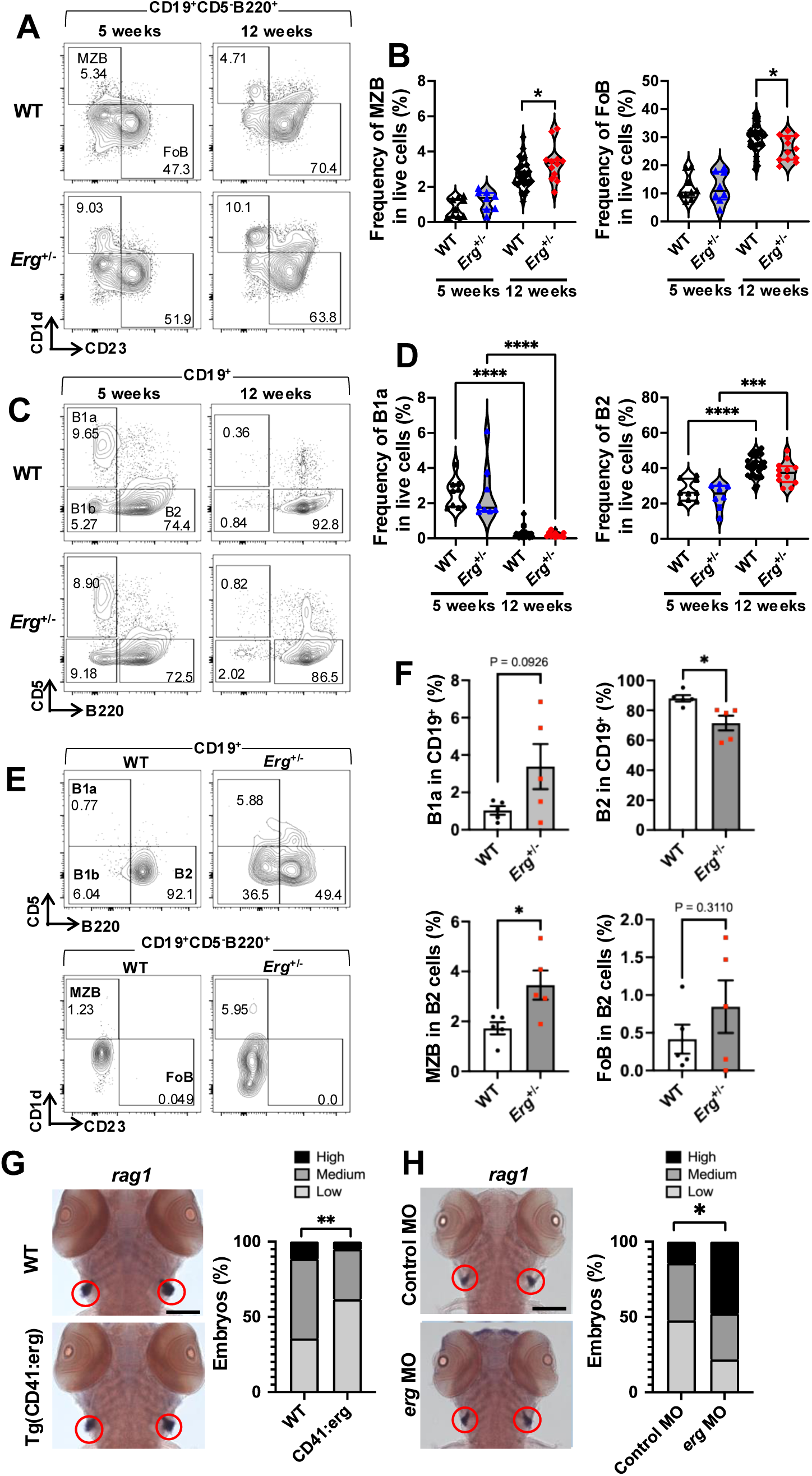
*Erg* regulates maturation of the lymphoid compartment. (A) Representative distributions of marginal zone B (MZB) and follicular B (FoB) cells in spleen (SP) from in 5- and 12-week-old WT and *Erg*^+/-^ mice. (B) Frequency of MZB and FoB in B2 cells 5- and 12-week-old SP (n = 8-28 per group). The results were pooled from four independent experiments. (C) Representative distributions of B1a, B1b, and B2 in 5- and 12-week-old SP. (D) Frequency of B1a and B2 cells in 5- and 12-week-old SP (n = 8-28 per group). The results were pooled from four independent experiments. (E) Representative distribution of B1a, B1b, B2, MZB, and FoB in OP9 culture derived from adult WT or *Erg*^+/-^ LSKs. (F) Frequency of B1a, B2, MZB, and FoB in OP9 derived from adult WT or *Erg*^+/-^ LSKs (n = 5 per group). The results were pooled from three independent experiments. (G) WISH for *rag1* at 5 dpf in thymus (red circle) in WT and Tg(CD41:*erg*) zebrafish and phenotypic distribution plot (WT, n = 87, Tg, n = 99). The results were pooled from four independent experiments. Scale bar, 100 µm. (H) WISH for *rag1* at 5 dpf in thymus (red circle) in control morpholino (MO) and *erg* MO zebrafish and phenotypic distribution plot (control MO, n = 22, erg MO, n = 23). The results were pooled from five independent experiments. Scale bar, 100 µm. Data are shown as mean ± SEM. Statistical tests were one-way ANOVA with Tukey’s multiple comparisons test (B, D, G), Student’s t-test (F), or Chi-square test (H-I). \**P* < 0.05; \*\**P* < 0.01; \*\*\**P* < 0.001; \*\*\*\**P* < 0.0001.

### Persistence of juvenile HSC phenotypes in *Erg* haploinsufficient adults

Juvenile and adult definitive HSCs differ in rates of self-renewal, transcriptional profiles, metabolic states, lineage biases, and surface marker expression^3,4,6–8,14,37–39^. At the phenotypic level, with aging, a subset of adult HSCs gains expression of cell surface CD41 as an indicator of MK potential via an alternative direct differentiation pathway^14,40^. We found that although there was no difference in CD41 expression on FL phenotypic long-term (LT)-HSCs, emergence of CD41^+^ LT-HSCs in the mature adult was impaired in *Erg*^+/-^ mice (Figure 4A-B, S2B). Consistent with a role for Erg in the emergence of this MK-primed subset of HSCs, among adult-biased HSPC ATAC peaks containing Erg binding motifs were several genes related to MK specification including *Pf4* and *Mpl* (Figure 4C, S4A). In orthogonal human datasets, we detected binding of ERG to human MK-associated loci (Figure S4B)^41^. LT-HSCs effectively and rapidly renew in the FL niche without exhaustion, and then at about four weeks following birth, self-renewal abruptly diminishes with LT-HSCs assuming a predominantly quiescent state in the adult BM^3^. By tracking the cell cycle state of Lineage^-^ Sca-1^+^ c-kit^+^ multipotent HSPCs (LSKs) and HSCs, we found that *Erg*^+/-^ LSKs/HSCs do not assume the scheduled quiescence in adulthood (Figure 4D-G). Self-renewal and quiescence of HSCs tracks with metabolic state – fetal HSCs show higher utilization of oxidative mitochondrial metabolism, while quiescent adult HSCs rely on glycolysis^38^. In alignment with our cell cycle data, we found that adult *Erg*^+/-^ LSKs/HSCs showed higher mitochondrial membrane polarization compared to littermates (Figure 4H-J). In support of these findings, morpholino knockdown of *erg* expression in the zebrafish CHT promotes HSC expansion (marked by *cmyb*; Figure 4K)^42^.

**Figure 4.**
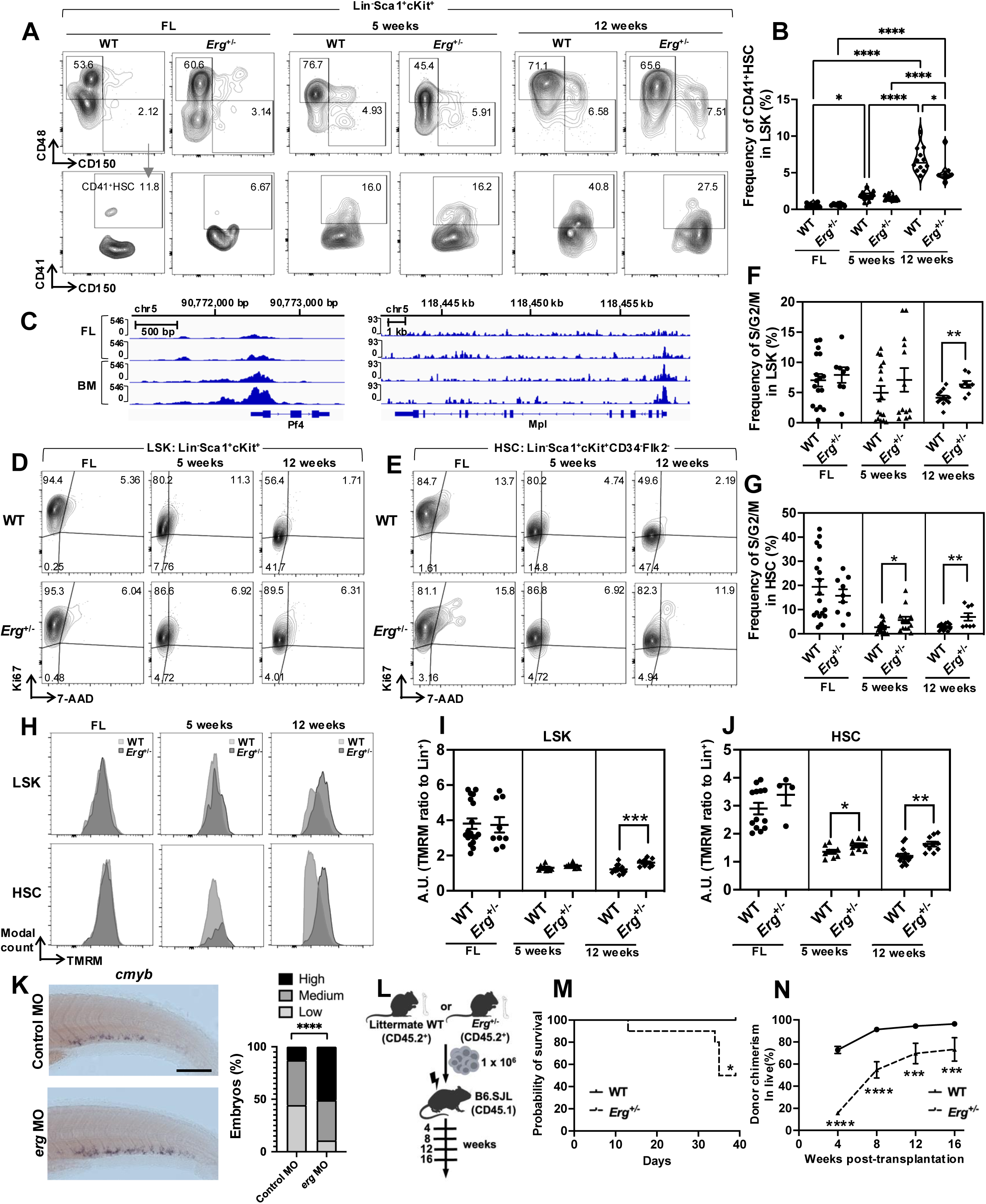
*Erg* regulates HSC developmental state. (A) Representative flow plots of CD41^+^HSCs in fetal (E14.5), 5-, and 12-week-old WT versus *Erg*^+/-^ mice. (B) Frequency of CD41^+^HSC in LSK from E14.5 FL, 5-, and 12-week-old BM (n = 8-14 per group). The results were pooled from six independent experiments. (C) ATAC-seq tracks at the *Pf4* and *Mpl* loci in FL HSC (n = 2) and adult BM HSC (n = 2)^14^. (D) Representative cell cycling assays in LSK from E14.5 FL, 5-, and 12-week-old BM. (E) Representative cell cycling assays in HSCs from E14.5 FL, 5-, and 12-week-old BM. (F) Frequency of S/G_2_/M phases in LSK from E14.5 FL, 5-, and 12-week-old BM (n = 8-17 per group). The results were pooled from eight independent experiments. (G) Frequency of S/G_2_/M phases in HSC from E14.5 FL, 5-, and 12-week-old BM (n = 8-18 per group). The results were pooled from eight independent experiments. (H) Representative histogram for mitochondrial membrane potential (𝚿) by TMRM stain in LSK and HSC from E14.5 FL, 5-, and 12-week-old BM. (I) Quantification of mean fluorescence intensity (MFI) of TMRM in LSK from E14.5 FL, 5-, and 12-week-old BM (n = 8-18 per group). The results were pooled from eight independent experiments. (J) Quantification of MFI of TMRM in HSC from E14.5 FL, 5-, and 12-week-old BM (n = 8-18 per group). The results were pooled from eight independent experiments. (K) WISH for *cmyb* at 3 dpf in CHT in control MO and *erg* MO zebrafish with phenotypic distribution plot (Control MO, n = 63, erg MO, n = 65). The results were pooled from four independent experiments. Scale bar, 100 µm. (L) Experimental design for non-competitive whole bone marrow transplantation. (M) Survival curve (n = 9-10 per group). Data is representative of two independent experiments. (N) Donor chimerism (CD45.2^+^) in peripheral blood (PB) over the course of the experiment (n = 9-10 per group). Data is representative of two independent experiments. Data are shown as mean ± SEM. Statistical tests were one-way ANOVA with Tukey’s multiple comparisons test (B), unpaired Student’s t test (F, G, I, J), Chi-square test (K), Fisher’s exact test (M), or two-way ANOVA with Sidak’s multiple comparisons test (N). \**P* < 0.05; \*\**P* < 0.01; \*\*\**P* < 0.001; \*\*\*\**P* < 0.0001.

Inappropriate activation and self-renewal of HSCs in the adult leads to replicative exhaustion^43,44^. *Erg*^+/-^ adults did not show significant expansion of the steady state HSC compartment and showed mild peripheral blood cytopenias suggestive of overall HSC dysfunction (Figure S4C-G, Table S1). Accordingly, transplantation of *Erg*^+/-^ BM into lethally irradiated recipients was not effective in rescuing survival compared to littermate BM (Figure 4L-N). Moreover, *Erg*^+/-^ HSCs did not perform as well as littermate HSCs in competitive transplantation (Figure S4H-J). Together, these results indicate that failure to downregulate rapid juvenile HSC self-renewal in the adult state in *Erg*^+/-^ adults is associated with replicative exhaustion.

### Persistence of fetal HSPC gene expression programs in *Erg*^+/-^ adults

To understand how Erg controls programs regulating the fetal to adult hematopoietic transition, we performed RNA sequencing (RNA-seq) on Lineage^-^ HSPCs of defined genotypes and ages. By RNA-seq, we identified transcripts differentially expressed between fetal and adult HSPCs, which represents the transcriptional program of the normal juvenile to adult transition (Figure 5A-C). In agreement with the hematopoietic phenotypes observed, we found that the transcriptional program of the juvenile to adult transition is impaired in *Erg*^+/-^ adult HSPCs with persistence of a portion of fetal patterns of gene expression postnatally (Figure 5A-C). Additionally, *Erg*^+/-^ HSPCs showed persistence of expression of fetal-biased TFs, including *Hmga2*, *Sall4*, and *Sox6*, suggesting that Erg plays a key role in downregulation of fetal HSPC TF networks (Figure 5D)^8,9,45,46^. Consistent with blunting of the age-related myeloid bias, *Erg*^+/-^ HSPCs showed diminished upregulation of the myeloid transcripts *Cpa3*, *Elane*, and *Mpo* (Figure 5E). Notably, we did not observe persistent expression of the juvenile HSPC transcripts *Lin28b*, *Igf2bp2*, and *Igf2bp3* in adulthood, consistent with *Erg* functioning downstream of these heterochronic factors (Figure S4J). However, by querying the overall developmental state of *Erg*^+/-^ adult HSPCs relative to wild-type adults using defined signatures of mouse and human fetal HSPCs, we found that adult *Erg*^+/-^ HSPCs retained overall juvenile HSPC transcriptional signatures (Figure 5F)^6,21,32^.

**Figure 5.**
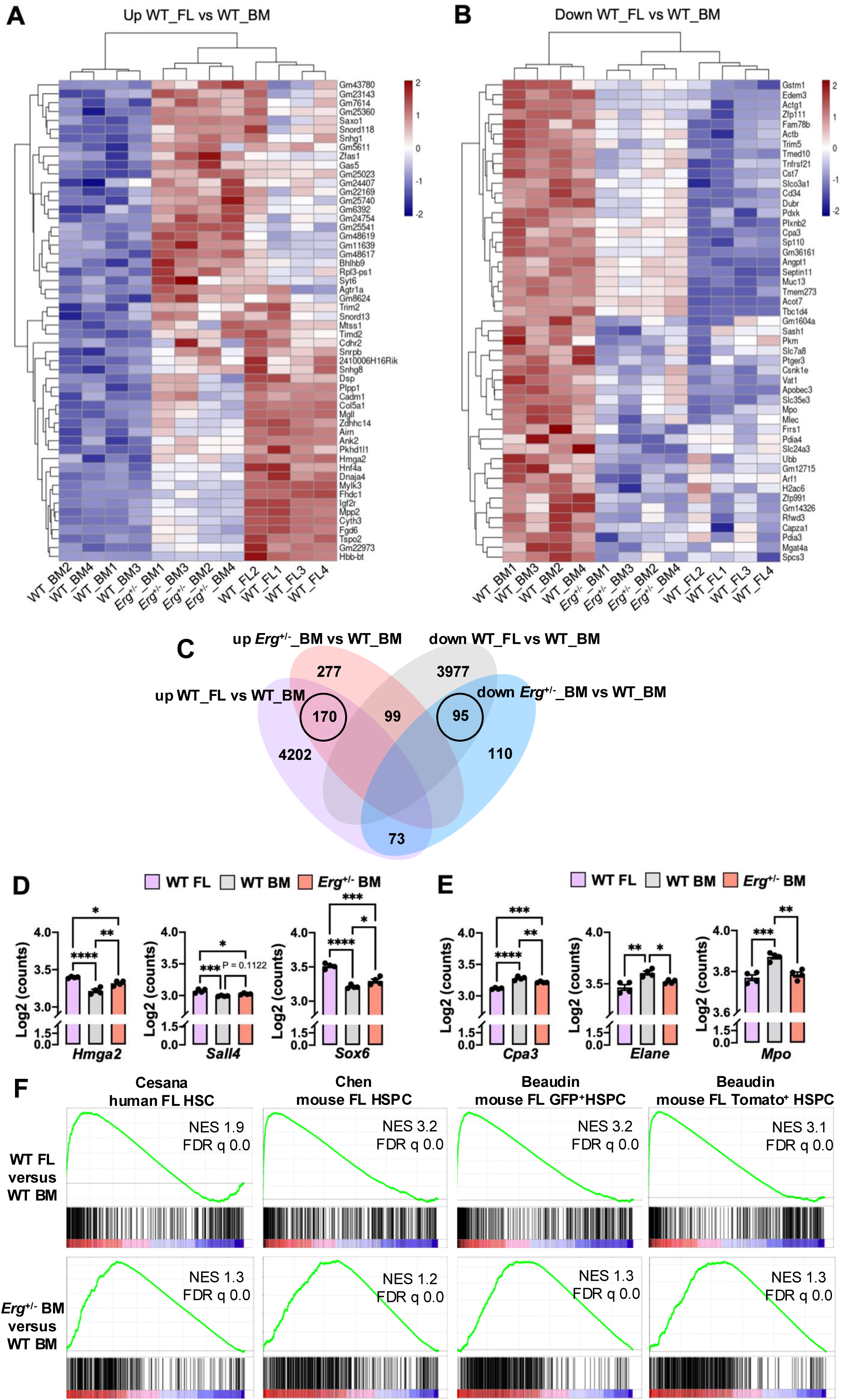
*Erg* regulates transcriptional and epigenetic programs in hematopoietic maturation. (A) Transcripts with fetal-biased expression also upregulated in *Erg*^+/-^ HSPCs relative to adult BM HSPCs. (B) Transcripts with adult-biased expression also downregulated in *Erg*^+/-^ HSPCs relative to adult BM HSPCs. (C) Venn diagram of the indicated comparisons, with regions corresponding to persistent fetal programs in *Erg*^+/-^ HSPCs circled. (D) Expression of key genes for transcription factors (*Hmga2*, *Sall4*, *Sox6*) in E14.5 FL, adult BM, *Erg*^+/-^BM HSPCs (n = 4 per group). (E) Expression of myeloid transcripts in E14.5 FL, adult BM, *Erg*^+/-^BM HSPCs (n = 4 per group). (F) GSEA of the indicated comparisons for the indicated human or mouse fetal HSPC signatures^6,21,32^.

### Mistimed *Hmga2* expression maintains fetal phenotypes

Hmga2 is a juvenile-biased TF that promotes HSC self-renewal and fetal hematopoietic output and functionally acts as a global regulator of chromatin structure and gene expression^8,9,47,48^. In human cell lines, we identified ERG binding at the human *HMGA2* locus including the promoter region, suggestive of regulation of *HMGA2* expression by ERG (Figure 6A)^41^. To query impact on chromatin structure, we performed chromatin immunoprecipitation with sequencing (ChIP-seq) for the active histone mark histone H3 lysine 27 acetylation (H3K27Ac). We identified discrete fetal and adult HSPC H3K27Ac peak profiles (Figure 6B). *Erg* haploinsufficiency in adult HSPCs resulted in partial persistence of fetal programs and incomplete implementation of adult programs in adult HSPCs (Figure 6B). Accordingly, by defining adult and fetal HSPC chromatin accessibility profiles based on ATAC-seq data, we observed that *Erg*^+/-^ impaired H3K27Ac localization to adult-specific accessible regions with persistence of this marker at fetal-specific accessible regions (Figure 6C^14^. Finally, consistent with Erg’s role in regulating age-specific HSPC TF networks, *Erg*^+/-^ HSPCs showed incomplete downregulation of fetal- and upregulation of adult-biased superenhancers (SEs; Figure 6D). Together, these studies show that full dose expression of *Erg* in the adult state is required for complete implementation of adult HSPC chromatin structure.

**Figure 6.**
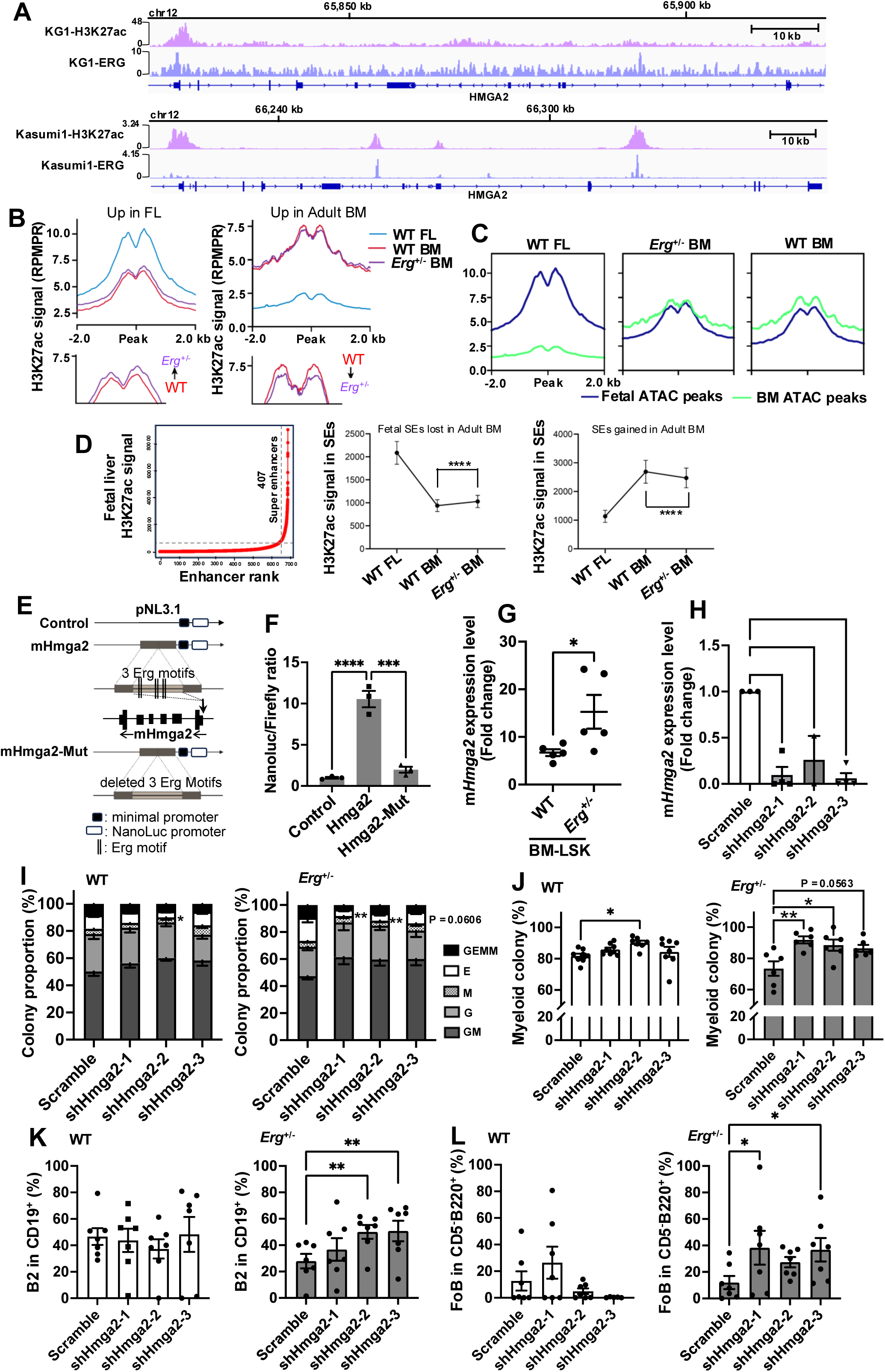
Persistent fetal phenotypes in *Erg*^+/-^ HSPCs are mediated by Hmga2. (A) KG1 (top tracks) or Kasumi-1 (bottom tracks) ChIP-seq data for H3K27Ac or ERG was visualized at the human *HMGA2* locus. (B) Representation of the indicated subsets of H3K27Ac ChIP-seq peaks (RPMPR; reads per million mapped peak reads). (C) Fetal- and adult-biased ATAC-seq peaks^14^ were identified and the distribution of these peaks visualized in the H3K27Ac ChIP-seq data of the indicated genotypes/age states. (D) Superenhancers (SEs) were identified and assigned to the WT FL or adult BM HSPCs with the relative H3K27Ac signal in these SEs quantified over each age/genotype. Data are shown as mean ± SEM. Statistical tests were one-way ANOVA with Tukey’s multiple comparisons test (D, E) and Wilcoxon rank test (H); \**P* < 0.05; \*\**P* < 0.01; \*\*\**P* < 0.001; \*\*\*\**P* < 0.0001. (E) Experimental design for reporter assay. ETS sites from a putative *Hmga2* enhancer were cloned into a luciferase expression vector (pNL3.1) upstream of a minimal promoter. An identical construct was generated with these sites mutated. (F) The indicated constructs (control: no enhancer cloned) were transfected into K562 cells and luciferase activity quantified (n = 3 technical replicates). Data is representative of three independent experiments. (G) qPCR showing levels of *Hmga2* expression in BM-LSK from WT or *Erg*^+/-^ mice (n = 3-5 per group). The results were pooled from four independent experiments. (H) Suppression of *Hmga2* expression in scramble, shRNA-*Hmga2* clones 1-3 (n = 2-4 per group). The results were pooled from three independent experiments. (I-J) CFU assay for transduced LSKs transduced with shRNAs against *Hmga2* from 12-week-old WT and *Erg*^+/-^ mice (n = 6-8 per group). The results were pooled from five independent experiments. Erythroid colonies were compared in (I), and total myeloid outcomes (G, M, GM) were compared in (J). (K-L) Frequency of B2 (K) or Fo (L) B-cells in OP9 culture from adult WT LSK and *Erg*^+/-^ LSKs expressing the indicated shRNAs (n = 7 per group). The results were pooled from four independent experiments). Data are shown as mean ± SEM. Statistical tests were one-way ANOVA with Tukey’s multiple comparisons test (F, H, I, J, K, L) and with unpaired Student’s t test (G); \**P* < 0.05; \*\**P* < 0.01; \*\*\**P* < 0.001; \*\*\*\**P* < 0.0001.

Persistent expression of *Hmga2* into adulthood could thus contribute to dysregulated HSC activation and juvenile hematopoietic phenotypes in *Erg*^+/-^ mice. We also identified several *Erg*/ETS family binding sites in putative enhancers adjacent to the mouse *Hmga2* locus, and we confirmed activity of these ETS sites in transcriptional activation using reporter assays (Figure 6E-F). We also confirmed elevated *Hmga2* expression in adult *Erg*^+/-^ LSKs by quantitative PCR (Figure 6G). To investigate a functional role for Hmga2 downstream of Erg, we generated small hairpin RNAs (shRNAs) targeting *Hmga2* and confirmed their ability to diminish *Hmga2* expression compared to a scramble control (Figure 6H). In myeloerythroid colony assays, while knockdown of *Hmga2* did not have significant effects in littermate adult HSPCs, knockdown of *Hmga2* in *Erg*^+/-^ HSPCs diminished erythroid output with skewing toward myeloid colony outcomes, consistent with blunting of this juvenile erythroid-skewed progenitor phenotype (Figure 6I-J)^9^. Furthermore, knockdown of *Hmga2* in adult *Erg*^+/-^ LSK HSPCs, but not in littermate control LSK HSPCs, promotes increased output of the adult B2 and Fo B-cell subpopulations (Figure 6K-L). Together, these findings support a model by which elevated expression of *Hmga2* at least in part mediates persistent juvenile hematopoietic phenotypes in *Erg* haploinsufficient adult HSPCs.

### *Erg* haploinsufficiency confers fetal-like resistance to transformation in HSPCs

Fetal HSPCs show relative resistance to oncogenic transformation compared to neonatal and adult HSPCs through mechanisms that are not fully understood but which appear to be independent of *Lin28b* expression^49,50^. To determine whether the persistent juvenile state of *Erg*^+/-^ HSPCs impacted susceptibility to transformation, we expressed the *MLL-AF9* oncogene in littermate control or *Erg*^+/-^ adult LSKs. We transplanted a defined quantity of transduced GFP^+^ LSKs into sublethally irradiated recipients and monitored survival (Figure 7A)^51^. We found that reduced dosage of *Erg* delayed onset of acute myeloid leukemia (AML) compared to littermates, akin to differences reported between oncogene induction fetal and postnatal cells (Figure 7B)^49,50^. *Erg*^+/-^ AMLs showed a more monocytic differentiation state (Figure 7C). These findings indicate that the persistent fetal state of *Erg*^+/-^ HSPCs impairs leukemogenesis.

**Figure 7.**
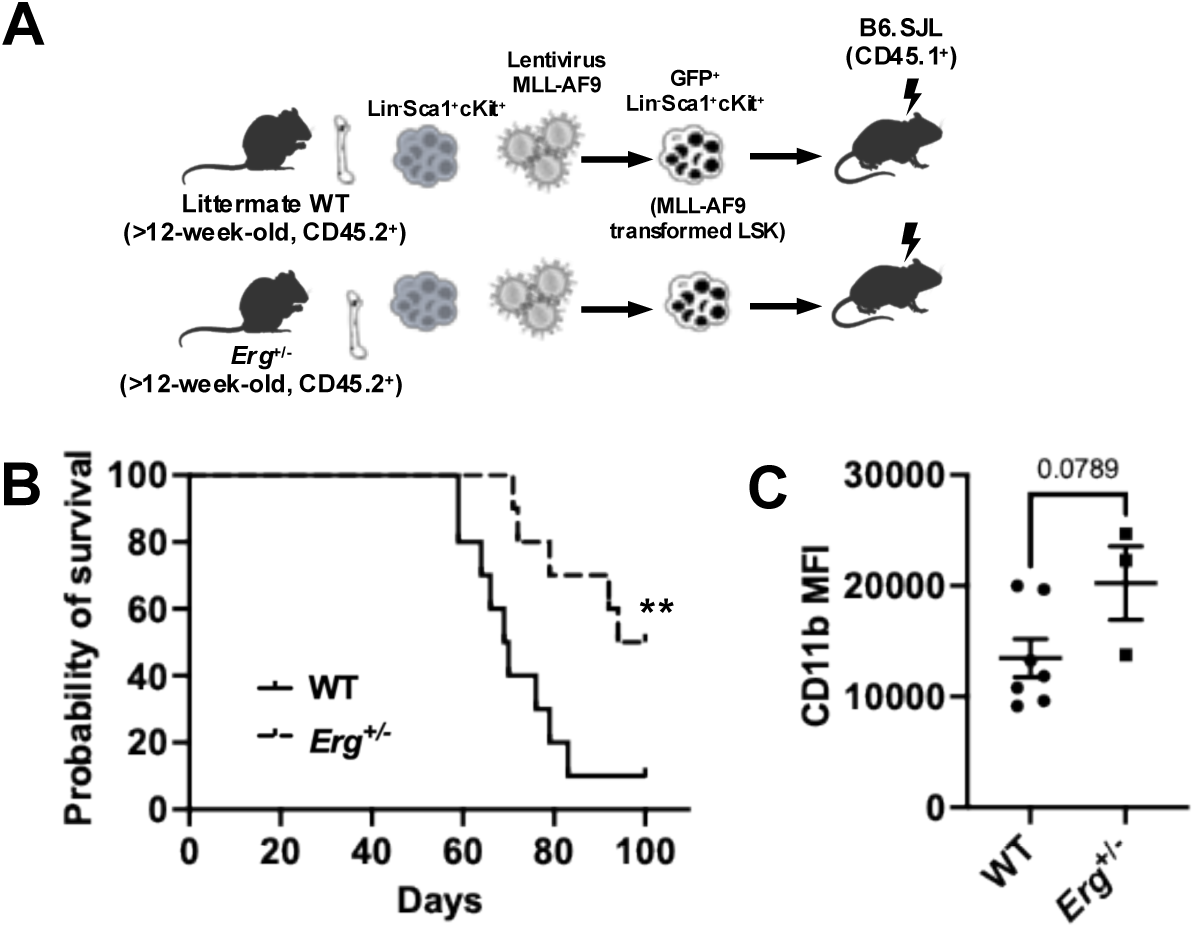
Impaired leukemogenesis in *Erg* haploinsufficiency. (A) Experimental schematic of *MLL-AF9* AML model. (B) Survival of mice transplanted with *MLL-AF9* leukemia (WT-*MLL-AF9*, n = 9; *Erg*^+/-^-*MLL-AF9*, n =10). (C) CD11b MFI from spleen at the endpoint (WT-*MLL-AF9*, n = 7; *Erg*^+/-^-*MLL-AF9*, n = 3). Statistical tests were Log-rank test (B) or unpaired Student’s t test (C); \*\**P* < 0.01.

## DISCUSSION

Here, we show that the master hematopoietic ETS family TF Erg is a heterochronic factor in definitive HSPCs with adult-biased expression necessary for completion of the juvenile to adult hematopoietic transition. These findings build on Erg’s known roles in early definitive HSC specification and in leukemogenesis^23–25,28,52^. ETS factors are central hematopoietic regulators, with stage-specific roles for ETV6 in HSC integrity posited previously, as suggested by its adult-biased expression as we observe^53,54^. Here, we implicate dynamic expression of Erg during hematopoietic aging in the implementation of the mature adult hematopoietic state.

Analysis of independent datasets and our own studies in mice demonstrate that temporally dynamic expression of *Erg*/*ERG*/*erg* within the hematopoietic system is evolutionarily conserved. Foundational studies in *Caenorhabditis elegans* first identified heterochronic factors including *lin-28* as being vital for the temporal coordination of developmental events, a paradigm which has been conserved in vertebrates in the hematopoietic system^55^. Detailed study has begun to uncover mechanisms by which Lin28b regulates phenotypic timing in vertebrates. Lin28 proteins bind RNA to regulate mRNA translation and the processing and maturation of microRNAs, predominantly of the *let-7* family^2,56^. Within the hematopoietic system, Lin28b is expressed in fetal HSPCs and is developmentally downregulated during the juvenile to adult transition, which enables maturation of *let-7* microRNAs that implement adult hematopoietic phenotypes^9,13^. Cbx2 is a *let-7* target transcript that is present in fetal HSPCs where Lin28b is expressed and *let-7* maturation is impaired^10^. Cbx2 is a PRC1 component that regulates the expression of hematopoietic TFs, including *Erg*^10^. Taken together, our findings support a model whereby developmental downregulation of *Lin28b* enables *let-7* maturation, decreasing Cbx2 levels and increasing expression of *Erg* during the juvenile to adult hematopoietic transition. As the direct *let-7* target *Hmga2* has also been implicated as an effector of Lin28b’s heterochronic control of HSPC function, our findings broaden these mechanisms to further explain the profound impact of Lin28b and *let-7* in regulating the developmental state of HSPCs^1,8–10^.

Several studies have indicated that *Erg* is required for maintenance of adult HSCs^22,27–29^. Adult HSCs must maintain quiescence, and dysregulated cycling due to either intrinsic loss of quiescence factors or exogenous stimuli such as inflammation leads to HSC exhaustion^43,57,58^. HSCs in the FL and in youth undergo rapid self-renewal without exhaustion to expand the developing hematopoietic system and then subsequently undergo a tightly programmed transition to quiescence in the BM at about four weeks of age in mice^3,59^. We find that loss of one *Erg* allele and consequent blunting of the programmed temporal increase in *Erg* expression results in persistent HSC cycling in adulthood with steady state cytopenias and impaired performance in transplantation assays. Interestingly, ectopic activation of Lin28b (which would be expected to repress *Erg* via Cbx2) drives HSC self-renewal within the adult BM without causing replicative exhaustion^8,10^. Erg therefore seems to play a role in maintaining appropriate adult HSC quiescence, while other factors downstream of Lin28b play positive roles in enabling maintenance of the HSC state during active cycling. Moreover, Lin28b-driven HSC self-renewal seems to be adaptive, as HSPCs expressing Lin28b are relatively resistant to oncogenic transformation, though this resistance to transformation may be due to downstream effects of Lin28b, i.e., secondary to an overall juvenile state^49,50^. This could be due to repression of the pro-oncogenic functions of Erg, as Erg expression is relatively lower in juvenile HSCs. Further study of Erg’s mechanistic regulation of HSC self-renewal and cell-intrinsic and -extrinsic control of HSC function in normal development and leukemia is therefore warranted.

The fetal to adult transition is characterized by adjustments in the proportions and qualities of mature blood cells produced which is underpinned by reprogramming at the level of HSPCs^5,9,12,13,60^. A prominent differentiation program that emerges upon transition to adult hematopoiesis is the alternative pathway of megakaryocyte production from CD41^+^ bone marrow HSCs^14^. Blunting temporal upregulation of *Erg* expression impairs the development of this primed HSC population and alternative MK differentiation pathway, as well as consequent downstream imbalances in MK subpopulations. As a mechanism, Erg seems to play a role in regulating the expression of genes that confer MK priming in adult CD41^+^ HSCs, likely also impacting production of adult-biased MkPs. Consistent with our findings, ectopic expression of *Erg* in fetal HSPCs results in CD41 expression and megakaryocytic output, which may partially explain its role in promoting megakaryoblastic leukemia in trisomy 21^26,61^.

Erg regulates other key aspects of hematopoietic aging. *Erg* haploinsufficiency results in persistent fetal-like erythroid skewing in the myeloerythroid progenitor compartment within the mature adult BM. In support of these findings, in human HSPCs, knockdown of *ERG* promotes erythroid output^41^. In the B-cell lineage, *Erg* has been shown to be crucial for full B-cell differentiation by regulating V(D)J recombination^31,62^. As Fo and MZ B-cells differ in B-cell receptor signaling, one possible mechanism by which temporally dynamic *Erg* expression regulates their specification is through V(D)J remodeling^63^. Thus, Erg appears to act at multiple levels of HSPC ontogeny to regulate self-renewal and fate choices during the aging of the hematopoietic system.

TF core regulatory circuitries (CRCs) are vital in defining cellular states^64^. We find that the transcriptional changes that occur during the juvenile to adult transition are partially abrogated in the setting of *Erg* haploinsufficiency. This includes impact on master TFs known to have stage specific effects in hematopoiesis. *Erg* haploinsufficiency also disrupts the programmed SE remodeling process of the juvenile to adult transition, consistent with direct impact on HSPC TF CRCs supported by our phenotypic data. TFs are central regulators of HSPC self-renewal and fate decisions^17^. Our results demonstrate the paradigm that TF expression can be dynamically regulated over aging to tailor hematopoietic output to transient physiologic demands.

While the TF Hmga2 is post-transcriptionally regulated by *let-7* to act as a heterochronic factor in HSPCs, we now demonstrate a relationship whereby Erg regulates *Hmga2* expression. *Erg* and *Hmga2* show inverse patterns of expression and appear to have opposing effects on the proliferation and activation of adult HSCs. Candidate fetal-specific HSPC enhancer sequences adjacent to *Hmga2* bear Erg/ETS bindings sites, indicating that Erg could act as a direct regulator of *Hmga2* expression, although other ETS factors likely also play roles in *Hmga2* expression^27,65^. In support of a linear relationship between these factors, downregulation of *Hmga2* specifically blunts juvenile-like hematopoietic programs maintained in *Erg*^+/-^ adult HSPCs. These findings provide a direct mechanism by which Erg interfaces with heterochronic regulators of HSC self-renewal during aging. The effective downregulation of *Lin28b* in *Erg*^+/-^ HSPCs would indicate that *Erg* functions downstream of Lin28b but is a regulator of *Hmga2* expression^10^. Thus, *Hmga2* appears to have dual levels of regulation during hematopoietic maturation – transcriptionally by ETS factors and post-transcriptionally by *let-7* microRNAs^66^. Further investigation is warranted to understand multilayered heterochronic regulation of Hmga2 by ETS factors.

From a disease-related perspective, activity of ERG is frequently perturbed in leukemia, and copy number gain of *ERG* on chromosome 21 has been implicated in the pathogenesis of Down syndrome-related and sporadic leukemia^23,25,26,67,68^. In pediatric leukemia, leukemogenic translocations often develop prenatally, but leukemia most often manifests after birth with several years of latency, consistent with the notion that fetal HSPCs are resistant to transformation, an effect putatively mediated by LIN28B or its effectors^50,69^. Our findings support a model wherein developmental upregulation of *ERG* during the fetal to postnatal transition plays a role in this differential susceptibility to leukemia. Moreover, aging is associated with an increased risk of myeloid malignancies due to temporal accumulation of somatic driver mutations in HSPCs^70,71^. However, given its pro-leukemic roles, our data raise the possibility that the age-related increase in *ERG* expression observed in human HSPCs plays a role in increased susceptibility to transformation by leukemic driver mutations with age.

Overall, our results demonstrate that although *Erg* is a globally required definitive hematopoietic TF, its expression and activity increase with aging and it is required for full implementation of the adult hematopoietic state, including the programmed transition to adult HSC quiescence. *Erg* haploinsufficiency results in persistence of hallmarks of juvenile hematopoiesis into adulthood; while *Erg* could be regulating these phenotypes independently, we also observe persistence of fetal HSPC gene expression programs in *Erg*^+/-^ adult hematopoiesis. Dynamic expression of *Erg* is regulated downstream of the master heterochronic Lin28b/*let-7* axis as a mechanism by which Lin28b exerts is widespread effects on blood formation. Further insight into the molecular mechanisms that specify programmed aging of definitive hematopoiesis will provide deeper understanding of blood diseases, many of which are skewed toward typical ages of onset.

## MATERIALS AND METHODS

### Mice and transplantation

All animal studies were approved by the Institutional Care and Use Committee at Boston Children’s Hospital. B6.Cg-Tg(Vav1-cre)1Graf/MdfJ (#035670) and *Erg*^tm1.1Iwamo^/J (#030988) mice were purchased from the Jackson Laboratory. The floxed *Erg* allele was deleted by targeting exon 6 with Vav1-cre mediated excision. Embryonic day 14.5 fetal liver (E14.5 FL), 5-week-old, and 12-week-old littermate wild type (WT) and Vav-Cre^+^;*Erg*^flox/+^(*Erg*^+/-^) mice were used. Transplant competitor cells and Sca-1 depleted rescue cells were isolated from CD45.1 (#002014, the Jackson Laboratory) mice. For non-competitive whole bone marrow transplantation, BM cells (1×10^6^) from 12-week-old WT and *Erg*^+/-^ mice were transplanted into lethally irradiated (975 rad) CD45.1 recipient mice. CD45.1 mice were used as recipients in competitive HSC transplantation as described. Peripheral blood analysis was performed at 4-week intervals post-transplantation. In the *MLL-AF9* model, LSK cells were isolated from adult WT or *Erg*^+/-^ mice and transduced with a retrovirus carrying the human MLL-AF9 cDNA. After one week of culture, GFP^+^ cells were sorted, and 4,000 GFP^+^ cells were injected via the tail vein sublethally irradiated (675 rad) CD45.1 recipient mice.

### Cell preparation from tissues

Mouse peripheral blood (PB) was obtained via retro-orbital plexus bleeding using heparinized micro-hematocrit capillary tubes. Bone marrow (BM) cells were collected from mouse hindlimb femur, pelvis, and tibia bones in sample medium containing DPBS, 2% FBS, 2 mM EDTA and kept on ice. Individual bones were dissociated using a mortar and pestle to ensure a single cell suspension, and the cell suspension was filtered through a 70 µm strainer. Spleens or thymi were collected in sample medium and kept on ice. Single cell suspensions were prepared by pushing the spleen or thymus tissue through a 70 µm strainer with a syringe plunger. The red blood cells from PB, BM, and SP were lysed by incubating the single cell suspension in ammonium-chloride-potassium (ACK) lysis buffer (155 mM NH_4_Cl, 10 mM KHCO_3_, 0.1 mM EDTA) and then washed twice with sample medium prior to analysis.

### Flow cytometry

#### Global population analysis

Details of the mouse antibodies and viability dyes used for each staining panel are shown in the Supplementary Figures. HSCs, progenitor cells, B cell subsets (B1a, B1b, B2, MZB, and FoB), and thymocytes were analyzed as previously described^72,73^. Cells were preincubated with LIVE/DEAD staining for 15 minutes, and then labeled with the appropriate cell-surface antibodies for 30 minutes at 4°C. LSRFortessa and FACSAria II were used for cell acquisition and cell sorting, respectively (BD Biosciences). All data analysis was performed using FlowJo software version 10.8.2 (FlowJo).

#### Cell cycle assay

After LIVE/DEAD and cell-surface antigen staining, cells were fixed and permeabilized with the BD Cytofix/Cytoperm Fixation/Permeabilization Solution Kit (BD Biosciences) and then stained with anti-Ki67 antibody (eBioScience) for 30 minutes at 4°C. After intracellular staining, cells were resuspended in sample medium containing 7-AAD (BioLegend) and incubated in the dark for 10 minutes before analysis.

#### Mitochondrial membrane potential assay

After staining for surface antigens, FL and BM cells were incubated with 2 nM tetramethylrhodamine, methyl ester, perchlorate (TMRM) in StemSPAN SFEM supplemented with 50 ng/mL SCF and TPO, and 50 µM verapamil for 60 minutes at 37°C followed by washing. Samples were resuspended in sample medium containing 2 nM TMRM. Data were collected on an LSRFortessa. Flow data analysis was performed using FlowJo.

### Reporter assays

To generate the Hmga2 reporter construct, a 226-nt with 3 ETS motifs (pNL3.1-mHmga2) or a 199-nt with 3 ETS motifs deleted (pNL3.1-mHmga2-Mut) was inserted into the pNL3.1 vector (#N1031, Promega). All three constructs were generated with Azenta/GENEWIZ. K562 cells were co-transfected with Lipofectamine 3000, pGL4.50 vector (Promega) and pNL3.1, pNL3.1-mHmga2, or pNL3.1-mHmga2-Mut. Twenty-four hours after transfection, cells were analyzed for luciferase activity using the Nano-Glo Dual-Luciferase Reporter Assay System (Promega) and signal was detected using SynergyTM NEO (BioTek). The normalized signal for firefly luciferase activity (NanoLuc luciferase activity/firefly luciferase activity) was calculated.

### Complete blood count

Peripheral Blood (100 µL) from mice was collected in lithium heparin-coated Microvette tubes (Sarstedt), then tubes were gently rotated for 10 min at room temperature. Complete blood cell count was analyzed using a Hemavet 950FS (Drew Scientific) according to the manufacturer’s instructions.

### OP9 co-culture

OP9 cells were plated at a density of 4×10^3^ cells/well in 96-well culture plates in OP9 medium (20% FBS in αMEM) 24 hours prior to OP9 co-culture initiation. BM cells were enriched by lineage depletion, and then live LSK or EGFP^+^ LSK cells were sorted by FACSAria II. Plate sorted 100 LSK cells were co-cultured with OP9 cells in αMEM supplemented with 10 ng/mL Flt3l, 20 ng/mL IL-7, 50 ng/mL SCF, and 20% FBS. The media was replaced every 3-4 days. After 10 days of culture, cells were analyzed by flow cytometry to define B-cell subtypes. Culture plates were maintained at 37°C, 5% CO_2_ and O_2_. Recombinant mouse cytokines were purchased from Peprotech.

### Quantitative PCR

RNA was purified with TRIzol Reagent, then cDNA synthesis was performed by using Maxima First Strand cDNA synthesis kit (Thermo). Real-time quantitative PCR was performed using QuantStudio 7 Flex real-time PCR system (Applied Biosystems) with Power SYBR Green PCR Master Mix (Thermo). Mouse *β-actin* primers were, F: 5’–ACGAGGCCCAGAGCAAGAGAGG–3’ and R: 5’–ACGCACCGATCCACACAGAGTA–3’. Mouse *Erg* primers were, F: 5’– GAGTGGGCGGTGAAAGAATA–3’ and R: 5’–TCAACGTCATCGGAAGTCAG–3’. Mouse *Hmga2* primers were, F: 5’–ATGGACTCATACACAGCAGCAG–3’ and R: 5’– AGGGAATATAGAGAGGAGAGAG–3’.

### Viral transduction

The lentiviral vector (pLV-EGFP:T2A:Puro-U6, VectorBuilder) was used to construct short hairpin RNA (shRNA) specific for mHmga2. The Hmga2-specific short hairpin RNA (shRNA, shHmga2) was 5’-TCGTTCAGAAGAAGCCTGCTC-3’ (shHmga2-1), 5’-AGACCCAGAGGAAGACCCAAA-3’ (shHmga2-2), 5’-GAAACTTATCAAGACGATTAA-3’ (shHmga2-3), and a non-specific scramble shRNA sequence was 5’-CCTAAGGTTAAGTCGCCCTCG-3’. HEK 293T cells were seeded at a concentration of 1×10^6^ cells/mL in a 150 mm culture dish with DMEM supplemented with 10% FBS. After 24 hours, cells were transfected with 9.75 ug total DNA per culture dish (3 scramble or shRNA for Hmga2: 2.25 pMD2G: 4.5 psPAX2). The retroviral vector pMIG-FLAG-MLL-AF9 (Addgene #71443) and pCL-Eco (Addgene #12371) were used for transfection to generate retrovirus^51^. Plasmids were mixed in 10 mL culture medium without FBS. Linear 40 kDa polyethylenimine (PEI, Polysciences) diluted to 8.0 mM in sterile water adjusted to pH 8.0 was used as transfection reagent. DNA-PEI complexes were incubated at room temperature for 20 minutes. Six hours after transfection, the plates were supplemented with fresh culture media containing 20% FBS. Cells were harvested 48 hours post-transfection at 37°C and the lentiviral supernatants were filtered through a 0.45 μm filter. Ultracentrifugation of lentivirus was performed with an SW28 rotor at 24,000 rpm for 2 hours. The obtained virus particle pellet was resuspended in Iscove’s modified Dulbecco’s medium (IMDM) and stored at −80°C. LSK cells were sorted by FACS into separate wells of a 96-well flat-bottom plate coated with RetroNectin (Takara), media containing 100 µL of IMDM supplemented with 10% FBS, 100 ng/mL Flt3l, 10 ng/mL IL-3, 100 ng/mL IL-6, 3 μg/mL polybrene, 100 ng/mL SCF, and 50 ng/mL TPO, and then spin-infected with lentivirus. Twenty-four hours after transduction, the medium was changed to fresh medium without polybrene. EGFP^+^ cells were sorted by FACS 72 hours after lentiviral transduction.

### Colony-forming assays

Following lentiviral transduction, EGFP^+^ LSK (Lin^-^Sca-1^+^cKit^+^) cells were sorted and plated in cytokine-supplemented complete methylcellulose medium (MethoCult GF M3434, STEMCELL Technologies). Erythroid colony-forming units (CFU-E), granulocyte colony-forming units (CFU-G), granulocyte-erythrocyte-macrophage-megakaryocyte colony-forming units (CFU-GEMM), granulocytes-macrophage colony-forming units (CFU-GM), and macrophage colony-forming units (CFU-M) were scored after 10-12 days according to the manufacturer’s protocol.

### Zebrafish

Zebrafish husbandry and experiments were performed according to approved IACUC protocols at Boston Children’s Hospital. Knockdown experiments were performed with morpholinos obtained from Gene Tools; clutches were split and injected with 15 ng of standard control MO (5’-CCTCTTACCTCAGTTACAATTTATA-3’) or a previously published translation blocking MO against zebrafish *erg* (erg-MO2, 5′-AGTGACACTCACTCTCTCTGAGGTA-3′)^74^. To generate the *Tg(cd41:erg; crybb1:TagBFP)* line, standard Gateway cloning techniques were performed. In brief, an LR reaction was used to combine the pCB24 pDEST crybb1:TagBFP backbone (Addgene plasmid #195987) with a p5E-CD41 promoter, a pME-erg zebrafish full-length cDNA (Horizon Discovery clone #ODR5433-202540699) and p3E-SV40 polyA sequence from the zebrafish Tol2 kit^75^. 50ng of purified DNA was coinjected with 30ng Tol2 transposase mRNA into one-cell stage AB(WT) embryos. Founders were screened by outcross to AB(WT) and sorted for individuals with blue eyes. All experiments were performed in the F2 offspring from F1 adults. For experiments, single *Tg(cd41:erg; crybb1:TagBFP)* fish were crossed to single AB(WT); the resulting clutches were split into BFP-controls and BFP+ siblings overexpressing erg under control of the CD41 promoter. At the desired timepoint of analysis, embryos were fixed in 4% PFA. Standard whole mount in situ protocols were performed to analyze lineage marker gene expression using previously published probes for *cmyb*, *mpo* and *rag1*. In situs were quantified by scoring all embryos within a stained clutch for relative distribution of staining intensities (typically high/medium/low), and the erg overexpressing embryos evaluated against their transgene-negative siblings (or erg morphants against standard control morphant controls) for shifted distribution by Chi-squared test. Results were depicted graphically in distribution plots, depicting scoring summary from >50 embryos from 2-4 experiments.

### ATAC-sequencing data analysis

scATAC-seq data was obtained from Gene Expression Omnibus with accession number GSE161724^14^. The sequencing data were processed to remove adapters and low-quality reads with Trimmomatic. Trimmed reads were aligned to the mouse reference genome (GRCm39) by BWA with default parameters. The fragment files were analysed using ArchR as a tile matrix with default parameters. Potential doublets were removed using filterDoublets function and pseudobulk replicates were generated using the addGroupCoverages function with default parameters. Bulk ATAC-seq data was obtained from Gene Expression Omnibus with accession numbers GSE161724^14^ and GSE119201^32^. Bulk ATAC-seq and pseudobulk files were processed for differential peak calling via the MACS2 algorithm and motif analysis using the HOMER algorithm on the Basepair platform.

### RNA-seq and data analysis

Lineage negative cells from WT FL (14.5), WT BM, and *Erg*^+/-^ BM, were negatively selected by biotinylated antibodies and streptavidin conjugated beads. RNA was purified with TRIzol Reagent and was treated with DNase I. Sequencing of whole RNA was performed at the Molecular Biology Core Facility at the Dana-Farber Cancer Institute. Libraries were prepared using SMARTer Stranded Total RNAseq v3 Pico Input Mammalian sample preparation kits from 5ng of purified total RNA according to the manufacturer’s protocol. The finished dsDNA libraries were quantified by Qubit fluorometer and Agilent TapeStation 4200. Uniquely dual indexed libraries were pooled in an equimolar ratio and shallowly sequenced on an Illumina MiSeq to further evaluate library quality and pool balance. The final pool was sequenced on an Illumina NovaSeq 6000 paired-end 150bp reads. Sequencing adapters and low-quality reads were removed with Trimmomatic. Trimmed reads were aligned to the mouse reference genome (GRCm39) by STAR with default parameters. Aligned reads were mapped to features using HTSeq, and differential expression analyses were performed using the DESeq2. Genes having less than 4 counts were excluded prior to statistical analysis, and differentially expressed genes (DEGs) were selected using as cutoffs the adjusted p-value < 0.05. Heatmaps were performed using pheatmap and list of DEGs was passed to enrichR and clusterProfiler for enrichment analyses. For gene set enrichment analysis (GSEA), the top 500 transcripts enriched in fetal liver versus adult marrow HSC/HSPCs in each referenced dataset were compiled into a signature that was used for GSEA for the indicated comparisons using our RNA-seq data^6,21,32^. For the Beaudin dataset, heterogeneous FL HSPCs are marked by either GFP or Tomato reporters, and differential expression versus adult marrow HSPCs is provided for both^6^.

The normalized expression of *ERG* in the published human single cell RNA-seq dataset was grouped into four age ranges based on the age of the donor (childhood 0-10 years, adolescence 11-20 years, adult 21-60 years, elderly 61+ years) and compared by the Wilcoxon rank-sum test^34^.

### ChIP-seq and data analysis

For each experiment, 10×10^6^ lineage negative cells were washed with PBS and flash frozen. Cells were then thawed and fixed with DSG and 1% formaldehyde, then digested with MNase for 15 minutes at 22°C and stopped with EGTA. Cells were then lysed with RIPA buffer, and chromatin was evaluated for digestion efficiency, achieving 50-70% mononucleosomes. Immunoprecipitation was carried out with anti-H3K27ac (Millipore, MABE647). After immunoprecipitation, samples were washed twice with RIPA buffer, twice with high salt (400mM NaCl) wash buffer, twice with LiCl (250mM) wash buffer, and twice with 150 mM NaCl buffer. Bead-bound chromatin was then end polished, followed by crosslink reversal and proteinase K digestion, and DNA purification with SPRI select beads. Libraries were prepared by ligating Illumina adaptors with barcoded index PCR. Each ChIP-seq sample was sequenced to a depth of 300 million or more paired-end reads with an Illumina NovaSeq 6000. Paired-end reads were aligned to human genome build mm10, using BWA version 0.7.17. The significance of peaks was tested using MACS3 (version 3.0.0a6, https://github.com/taoliu/MACS) with the -B option, a regular TF-binding mode. Peak artifacts were removed (reference locations “black-listed” by the ENCODE consortium https://sites.google.com/site/anshulkundaje/projects/blacklists, ENCFF356LFX). Super enhancers were called using the ROSE algorithm (v2). H3K27ac signal was normalized as Reads Per Million Peak Reads. Bedtools coverage was used to calculate the number of reads in regions of interest. Density profiles were plotted via the Deeptools package.

Datasets available in Gene Expression Omnibus (GEO) of ERG ChIP-seq analyzed were GSE167163 and GSE158794^41^.

## DATA AVAILABILITY STATEMENT

Requests for data should be addressed to. RNA-seq and ChIP-seq data are available on GEO with accession number GSE269184.

## ACKNOWLEDGEMENTS

This work was supported by NIDDK R01DK134515, R03DK126729 (to R.G.R.) and K01DK129409 (to W.W.S.), Coordination for the Improvement of Higher Education Personnel, Serrapilheira Institute and Research and Innovation Support Foundation of the State of Santa Catarina (to E.L.R), and DOD Convergent Science Virtual Cancer Center Scholar Award W81XWH-21-1-0298 (to B.E.G.). We thank the staff of the Flow Cytometry Core, Animal Resources Children’s Hospital at the Boston Children’s Hospital, and Dr. Elizabeth Molnar for zebrafish husbandry and support.

## AUTHOR CONTRIBUTIONS

M.T.Y., W.W.S., D.W., B.B., P.C., D.C., J.G., S.G., and G.R-L. performed research; M.T.Y., W.W.S., E.L.R., H.L. and B.G. analyzed data; B.G., T.E.N., and R.G.R. designed research, R.G.R. wrote the manuscript.

## REFERENCES

1 Rowe, R. G., Mandelbaum, J., Zon, L. I. & Daley, G. Q. Engineering Hematopoietic Stem Cells: Lessons from Development. Cell Stem Cell 18, 707–720 (2016). 10.1016/j.stem.2016.05.016

2 Basak, A. et al. Control of human hemoglobin switching by LIN28B-mediated regulation of BCL11A translation. Nat Genet (2020). 10.1038/s41588-019-0568-7

3 Bowie, M. B. et al. Identification of a new intrinsically timed developmental checkpoint that reprograms key hematopoietic stem cell properties. Proc Natl Acad Sci U S A 104, 5878–5882 (2007). 10.1073/pnas.0700460104

4 Notta, F. et al. Distinct routes of lineage development reshape the human blood hierarchy across ontogeny. Science 351, aab2116 (2016). 10.1126/science.aab2116

5 Yuan, J., Nguyen, C. K., Liu, X., Kanellopoulou, C. & Muljo, S. A. Lin28b reprograms adult bone marrow hematopoietic progenitors to mediate fetal-like lymphopoiesis. Science 335, 1195–1200 (2012). 10.1126/science.1216557

6 Beaudin, A. E. et al. A Transient Developmental Hematopoietic Stem Cell Gives Rise to Innate-like B and T Cells. Cell Stem Cell 19, 768–783 (2016). 10.1016/j.stem.2016.08.013

7 Li, Y. et al. Single-Cell Analysis of Neonatal HSC Ontogeny Reveals Gradual and Uncoordinated Transcriptional Reprogramming that Begins before Birth. Cell Stem Cell 27, 732–747 e737 (2020). 10.1016/j.stem.2020.08.001

8 Copley, M. R. et al. The Lin28b-let-7-Hmga2 axis determines the higher self-renewal potential of fetal haematopoietic stem cells. Nat Cell Biol 15, 916–925 (2013). 10.1038/ncb2783

9 Rowe, R. G. et al. Developmental regulation of myeloerythroid progenitor function by the Lin28b-let-7-Hmga2 axis. J Exp Med 213, 1497–1512 (2016). 10.1084/jem.20151912

10 Wang, D. et al. Developmental maturation of the hematopoietic system controlled by a Lin28b-let-7-Cbx2 axis. Cell Rep 39, 110587 (2022). 10.1016/j.celrep.2022.110587

11 Shyh-Chang, N. & Daley, G. Q. Lin28: primal regulator of growth and metabolism in stem cells. Cell Stem Cell 12, 395–406 (2013). 10.1016/j.stem.2013.03.005

12 Lee, Y. T. et al. LIN28B-mediated expression of fetal hemoglobin and production of fetal-like erythrocytes from adult human erythroblasts ex vivo. Blood 122, 1034–1041 (2013). 10.1182/blood-2012-12-472308

13 Zhou, Y. et al. Lin28b promotes fetal B lymphopoiesis through the transcription factor Arid3a. J Exp Med 212, 569–580 (2015). 10.1084/jem.20141510

14 Kristiansen, T. A. et al. Developmental cues license megakaryocyte priming in murine hematopoietic stem cells. Blood Adv 6, 6228–6241 (2022). 10.1182/bloodadvances.2021006861

15 Stolla, M. C. et al. Lin28b regulates age-dependent differences in murine platelet function. Blood Adv 3, 72–82 (2019). 10.1182/bloodadvances.2018020859

16 Wang, L. D. et al. The role of Lin28b in myeloid and mast cell differentiation and mast cell malignancy. Leukemia 29, 1320–1330 (2015). 10.1038/leu.2015.19

17 Orkin, S. H. Transcription factors and hematopoietic development. J Biol Chem 270, 4955–4958 (1995). 10.1074/jbc.270.10.4955

18 de Bruijn, M. & Dzierzak, E. Runx transcription factors in the development and function of the definitive hematopoietic system. Blood 129, 2061–2069 (2017). 10.1182/blood-2016-12-689109

19 Look, A. T. Oncogenic transcription factors in the human acute leukemias. Science 278, 1059–1064 (1997). 10.1126/science.278.5340.1059

20 Kim, I., Saunders, T. L. & Morrison, S. J. Sox17 dependence distinguishes the transcriptional regulation of fetal from adult hematopoietic stem cells. Cell 130, 470–483 (2007). 10.1016/j.cell.2007.06.011

21 Cesana, M. et al. A CLK3-HMGA2 Alternative Splicing Axis Impacts Human Hematopoietic Stem Cell Molecular Identity throughout Development. Cell Stem Cell 22, 575–588 e577 (2018). 10.1016/j.stem.2018.03.012

22 Loughran, S. J. et al. The transcription factor Erg is essential for definitive hematopoiesis and the function of adult hematopoietic stem cells. Nat Immunol 9, 810–819 (2008). 10.1038/ni.1617

23 Tsuzuki, S., Taguchi, O. & Seto, M. Promotion and maintenance of leukemia by ERG. Blood 117, 3858–3868 (2011). 10.1182/blood-2010-11-320515

24 Qian, M. et al. Novel susceptibility variants at the ERG locus for childhood acute lymphoblastic leukemia in Hispanics. Blood 133, 724–729 (2019). 10.1182/blood-2018-07-862946

25 de Smith, A. J. et al. Heritable variation at the chromosome 21 gene ERG is associated with acute lymphoblastic leukemia risk in children with and without Down syndrome. Leukemia 33, 2746–2751 (2019). 10.1038/s41375-019-0514-9

26 Salek-Ardakani, S. et al. ERG is a megakaryocytic oncogene. Cancer Res 69, 4665–4673 (2009). 10.1158/0008-5472.CAN-09-0075

27 Knudsen, K. J. et al. ERG promotes the maintenance of hematopoietic stem cells by restricting their differentiation. Genes Dev 29, 1915–1929 (2015). 10.1101/gad.268409.115

28 Taoudi, S. et al. ERG dependence distinguishes developmental control of hematopoietic stem cell maintenance from hematopoietic specification. Genes Dev 25, 251–262 (2011). 10.1101/gad.2009211

29 Xie, Y. et al. Reduced Erg Dosage Impairs Survival of Hematopoietic Stem and Progenitor Cells. Stem Cells 35, 1773–1785 (2017). 10.1002/stem.2627

30 Ng, A. P. et al. Erg is required for self-renewal of hematopoietic stem cells during stress hematopoiesis in mice. Blood 118, 2454–2461 (2011). 10.1182/blood-2011-03-344739

31 Ng, A. P. et al. An Erg-driven transcriptional program controls B cell lymphopoiesis. Nat Commun 11, 3013 (2020). 10.1038/s41467-020-16828-y

32 Chen, C. et al. Spatial Genome Re-organization between Fetal and Adult Hematopoietic Stem Cells. Cell Rep 29, 4200–4211 e4207 (2019). 10.1016/j.celrep.2019.11.065

33 Xue, Y. et al. A 3D Atlas of Hematopoietic Stem and Progenitor Cell Expansion by Multi-dimensional RNA-Seq Analysis. Cell Rep 27, 1567–1578 e1565 (2019). 10.1016/j.celrep.2019.04.030

34 Zhang, Y. et al. Temporal molecular program of human hematopoietic stem and progenitor cells after birth. Dev Cell 57, 2745–2760 e2746 (2022). 10.1016/j.devcel.2022.11.013

35 Ohta, Y. et al. Articular cartilage endurance and resistance to osteoarthritic changes require transcription factor Erg. Arthritis Rheumatol 67, 2679–2690 (2015). 10.1002/art.39243

36 MacKinney, A. A., Jr. Effect of aging on the peripheral blood lymphocyte count. J Gerontol 33, 213–216 (1978). 10.1093/geronj/33.2.213

37 Rebel, V. I., Miller, C. L., Eaves, C. J. & Lansdorp, P. M. The repopulation potential of fetal liver hematopoietic stem cells in mice exceeds that of their liver adult bone marrow counterparts. Blood 87, 3500–3507 (1996).

38 Nakamura-Ishizu, A., Ito, K. & Suda, T. Hematopoietic Stem Cell Metabolism during Development and Aging. Dev Cell 54, 239–255 (2020). 10.1016/j.devcel.2020.06.029

39 Morrison, S. J., Hemmati, H. D., Wandycz, A. M. & Weissman, I. L. The purification and characterization of fetal liver hematopoietic stem cells. Proc Natl Acad Sci U S A 92, 10302–10306 (1995). 10.1073/pnas.92.22.10302

40 Sanjuan-Pla, A. et al. Platelet-biased stem cells reside at the apex of the haematopoietic stem-cell hierarchy. Nature 502, 232–236 (2013). 10.1038/nature12495

41 Thoms, J. A. I. et al. Disruption of a GATA2-TAL1-ERG regulatory circuit promotes erythroid transition in healthy and leukemic stem cells. Blood 138, 1441–1455 (2021). 10.1182/blood.2020009707

42 North, T. E. et al. Prostaglandin E2 regulates vertebrate haematopoietic stem cell homeostasis. Nature 447, 1007–1011 (2007). 10.1038/nature05883

43 Yang, Y. et al. The histone lysine acetyltransferase HBO1 (KAT7) regulates hematopoietic stem cell quiescence and self-renewal. Blood 139, 845–858 (2022). 10.1182/blood.2021013954

44 Kamminga, L. M. et al. The Polycomb group gene Ezh2 prevents hematopoietic stem cell exhaustion. Blood 107, 2170–2179 (2006). 10.1182/blood-2005-09-3585

45 Xu, J. et al. Transcriptional silencing of gamma-globin by BCL11A involves long-range interactions and cooperation with SOX6. Genes Dev 24, 783–798 (2010). 10.1101/gad.1897310

46 Huang, P. et al. HIC2 controls developmental hemoglobin switching by repressing BCL11A transcription. Nat Genet 54, 1417–1426 (2022). 10.1038/s41588-022-01152-6

47 Kuwayama, N. et al. HMGA2 directly mediates chromatin condensation in association with neuronal fate regulation. Nat Commun 14, 6420 (2023). 10.1038/s41467-023-42094-9

48 Zhang, Q. & Wang, Y. HMG modifications and nuclear function. Biochim Biophys Acta 1799, 28-36 (2010). 10.1016/j.bbagrm.2009.11.009

49 Li, Y. et al. LIN28B promotes differentiation of fully transformed AML cells but is dispensable for fetal leukemia suppression. Leukemia 38, 648–651 (2024). 10.1038/s41375-024-02167-0

50 Okeyo-Owuor, T. et al. The efficiency of murine MLL-ENL-driven leukemia initiation changes with age and peaks during neonatal development. Blood Adv 3, 2388–2399 (2019). 10.1182/bloodadvances.2019000554

51 Rowe, R. G. et al. The developmental stage of the hematopoietic niche regulates lineage in MLL-rearranged leukemia. J Exp Med 216, 527–538 (2019). 10.1084/jem.20181765

52 Carmichael, C. L. et al. Hematopoietic overexpression of the transcription factor Erg induces lymphoid and erythro-megakaryocytic leukemia. Proc Natl Acad Sci U S A 109, 15437–15442 (2012). 10.1073/pnas.1213454109

53 Wang, L. C. et al. The TEL/ETV6 gene is required specifically for hematopoiesis in the bone marrow. Genes Dev 12, 2392–2402 (1998). 10.1101/gad.12.15.2392

54 Hock, H. et al. Tel/Etv6 is an essential and selective regulator of adult hematopoietic stem cell survival. Genes Dev 18, 2336–2341 (2004). 10.1101/gad.1239604

55 Ambros, V. & Horvitz, H. R. Heterochronic mutants of the nematode Caenorhabditis elegans. Science 226, 409–416 (1984). 10.1126/science.6494891

56 Viswanathan, S. R., Daley, G. Q. & Gregory, R. I. Selective blockade of microRNA processing by Lin28. Science 320, 97–100 (2008). 10.1126/science.1154040

57 Caiado, F., Pietras, E. M. & Manz, M. G. Inflammation as a regulator of hematopoietic stem cell function in disease, aging, and clonal selection. J Exp Med 218 (2021). 10.1084/jem.20201541

58 Bogeska, R. et al. Inflammatory exposure drives long-lived impairment of hematopoietic stem cell self-renewal activity and accelerated aging. Cell Stem Cell 29, 1273–1284 e1278 (2022). 10.1016/j.stem.2022.06.012

59 Bowie, M. B. et al. Hematopoietic stem cells proliferate until after birth and show a reversible phase-specific engraftment defect. J Clin Invest 116, 2808–2816 (2006). 10.1172/JCI28310

60 de Vasconcellos, J. F. et al. IGF2BP1 overexpression causes fetal-like hemoglobin expression patterns in cultured human adult erythroblasts. Proc Natl Acad Sci U S A 114, E5664–E5672 (2017). 10.1073/pnas.1609552114

61 Stankiewicz, M. J. & Crispino, J. D. ETS2 and ERG promote megakaryopoiesis and synergize with alterations in GATA-1 to immortalize hematopoietic progenitor cells. Blood 113, 3337–3347 (2009). 10.1182/blood-2008-08-174813

62 Sondergaard, E. et al. ERG Controls B Cell Development by Promoting Igh V-to-DJ Recombination. Cell Rep 29, 2756–2769 e2756 (2019). 10.1016/j.celrep.2019.10.098

63 Cerutti, A., Cols, M. & Puga, I. Marginal zone B cells: virtues of innate-like antibody-producing lymphocytes. Nat Rev Immunol 13, 118–132 (2013). 10.1038/nri3383

64 Saint-Andre, V. et al. Models of human core transcriptional regulatory circuitries. Genome Res 26, 385–396 (2016). 10.1101/gr.197590.115

65 Dryden, N. H. et al. The transcription factor Erg controls endothelial cell quiescence by repressing activity of nuclear factor (NF)-kappaB p65. J Biol Chem 287, 12331–12342 (2012). 10.1074/jbc.M112.346791

66 Lee, Y. S. & Dutta, A. The tumor suppressor microRNA let-7 represses the HMGA2 oncogene. Genes Dev 21, 1025–1030 (2007). 10.1101/gad.1540407

67 Schmoellerl, J. et al. EVI1 drives leukemogenesis through aberrant ERG activation. Blood 141, 453–466 (2023). 10.1182/blood.2022016592

68 Gao, Q. et al. The genomic landscape of acute lymphoblastic leukemia with intrachromosomal amplification of chromosome 21. Blood 142, 711–723 (2023). 10.1182/blood.2022019094

69 Eldeeb, M. et al. A fetal tumor suppressor axis abrogates MLL-fusion-driven acute myeloid leukemia. Cell Rep 42, 112099 (2023). 10.1016/j.celrep.2023.112099

70 Jaiswal, S. & Ebert, B. L. Clonal hematopoiesis in human aging and disease. Science 366 (2019). 10.1126/science.aan4673

71 Ogawa, S. Genetics of MDS. Blood 133, 1049–1059 (2019). 10.1182/blood-2018-10-844621

72 Beerman, I. et al. Functionally distinct hematopoietic stem cells modulate hematopoietic lineage potential during aging by a mechanism of clonal expansion. Proc Natl Acad Sci U S A 107, 5465–5470 (2010). 10.1073/pnas.1000834107

73 Tanaka-Yano, M. et al. Tristetraprolin overexpression drives hematopoietic changes in young and middle-aged mice generating dominant mitigating effects on induced inflammation in murine models. Geroscience 46, 1271–1284 (2024). 10.1007/s11357-023-00879-2

74 Liu, F. & Patient, R. Genome-wide analysis of the zebrafish ETS family identifies three genes required for hemangioblast differentiation or angiogenesis. Circ Res 103, 1147–1154 (2008). 10.1161/CIRCRESAHA.108.179713

75 Kwan, K. M. et al. The Tol2kit: a multisite gateway-based construction kit for Tol2 transposon transgenesis constructs. Dev Dyn 236, 3088–3099 (2007). 10.1002/dvdy.21343

76 Heinz, S. et al. Simple combinations of lineage-determining transcription factors prime cis-regulatory elements required for macrophage and B cell identities. Mol Cell 38, 576–589 (2010). 10.1016/j.molcel.2010.05.004

