## Supplemental Figures 1-4, Supplemental Table 1 for "Dynamic activity of *Erg* promotes aging of the hematopoietic system"

**Yano_FigS1**

**A**

| Cell type | Fetal | Adult |
|---|---|---|
| HSC | Cell cycle↑<br>Mitochondrial membrane potential (ΔΨm) ↑ | Cell cycle↓<br>ΔΨm↓ |
| Myeloid Progenitors | MEP/GMP ratio ↑<br>MkP↓ | MEP/GMP ratio ↓<br>MkP↑ |
| Lymphoid Progenitors | CLP↑ | CLP↓ |
| B cells | B1a and MZB | B2 and FoB |
| T cells | $\gamma\delta$T | $\alpha\beta$T |

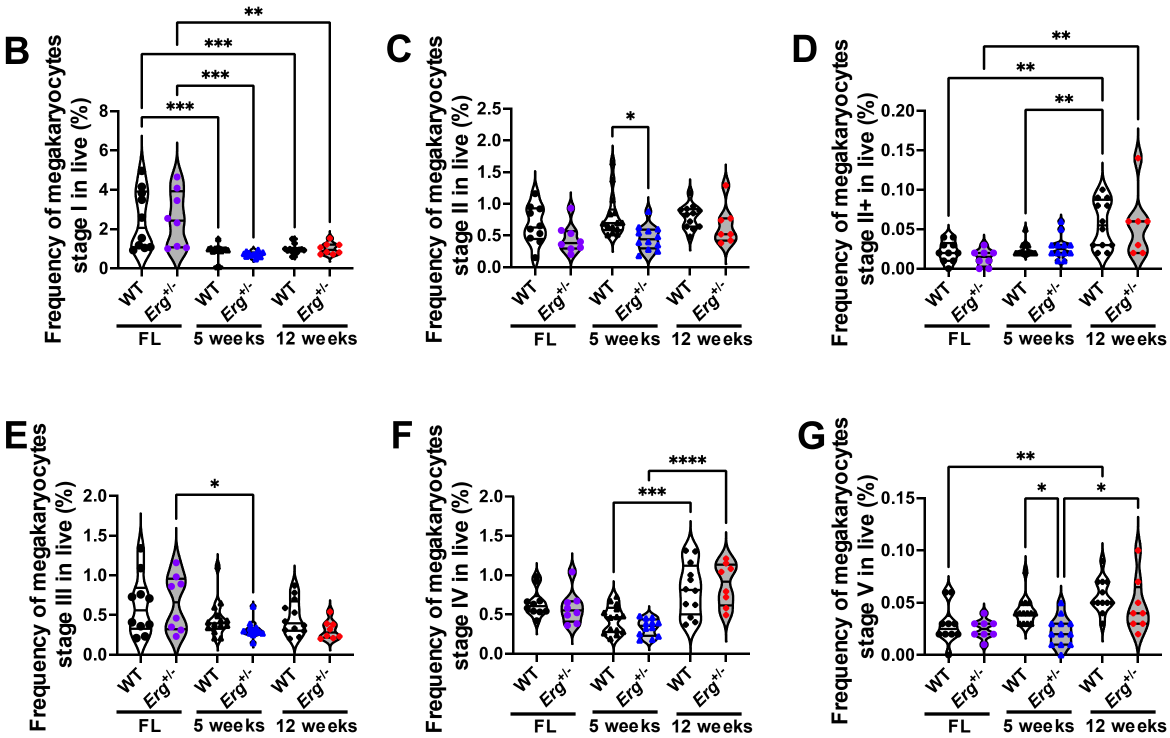

**A**

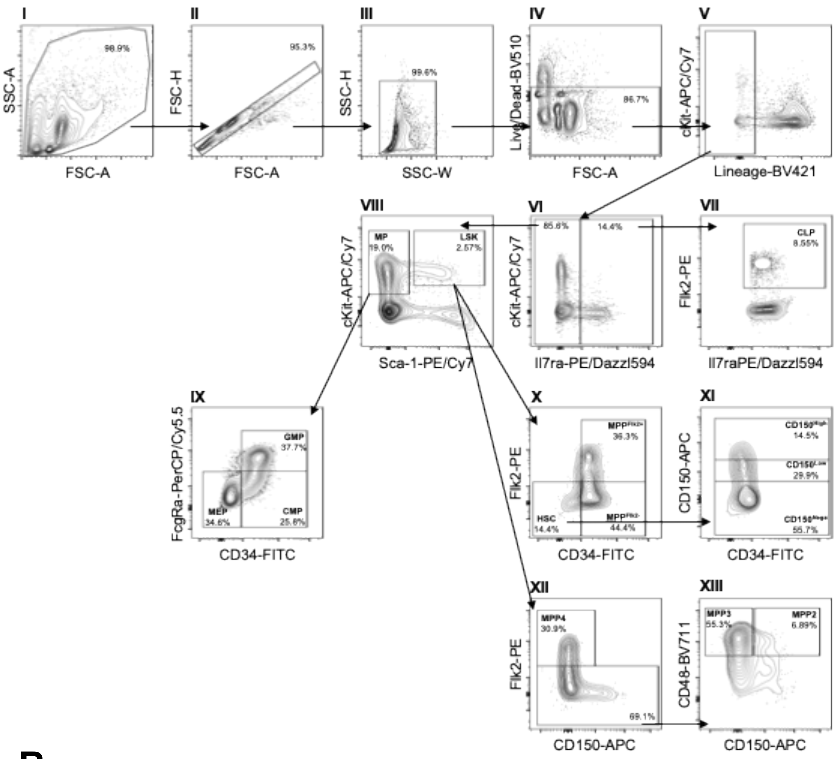


| Cell type | Immunophenotype |
|---|---|
| CLP | Lin$^-$ IL-7Ra$^+$ Flk2$^+$ |
| MP | Lin$^-$ IL-7Ra$^-$ Sca1$^+$ cKit$^-$ |
| CMP | Lin$^-$ IL-7Ra$^-$ Sca1$^+$ cKit$^-$ CD34$^+$ FcγRa$^-$ |
| GMP | Lin$^-$ IL-7Ra$^-$ Sca1$^+$ cKit$^-$ CD34$^+$ FcγRa$^+$ |
| MEP | Lin$^-$ IL-7Ra$^-$ Sca1$^+$ cKit$^-$ CD34$^-$ FcγRa$^-$ |
| LSK | Lin$^-$ IL-7Ra$^-$ Sca1$^+$ cKit$^+$ |
| HSC | Lin$^-$ IL-7Ra$^-$ Sca1$^+$ cKit$^+$ CD34$^-$ |
| MPP$^{Flk2+}$ | Lin$^-$ IL-7Ra$^-$ Sca1$^+$ cKit$^+$ CD34$^+$ Flk2$^+$ |
| MPP$^{Flk2-}$ | Lin$^-$ IL-7Ra$^-$ Sca1$^+$ cKit$^+$ CD34$^+$ Flk2$^-$ |
| CD150$^{Nega}$ | Lin$^-$ IL-7Ra$^-$ Sca1$^+$ cKit$^+$ CD34$^-$ CD150$^{Nega}$ |
| CD150$^{Low}$ | Lin$^-$ IL-7Ra$^-$ Sca1$^+$ cKit$^+$ CD34$^-$ CD150$^{Low}$ |
| CD150$^{High}$ | Lin$^-$ IL-7Ra$^-$ Sca1$^+$ cKit$^+$ CD34$^-$ CD150$^{High}$ |
| MPP2 | Lin$^-$ IL-7Ra$^-$ Sca1$^+$ cKit$^+$ Flk2$^-$ CD48$^+$ CD150$^+$ |
| MPP3 | Lin$^-$ IL-7Ra$^-$ Sca1$^+$ cKit$^+$ Flk2$^-$ CD48$^+$ CD150$^-$ |
| MPP4 | Lin$^-$ IL-7Ra$^-$ Sca1$^+$ cKit$^+$ Flk2$^+$ |

Lineage: B220, CD11b, CD3, Gr1, Ter119
(CD11b was excluded for fetal liver samples)

**B**

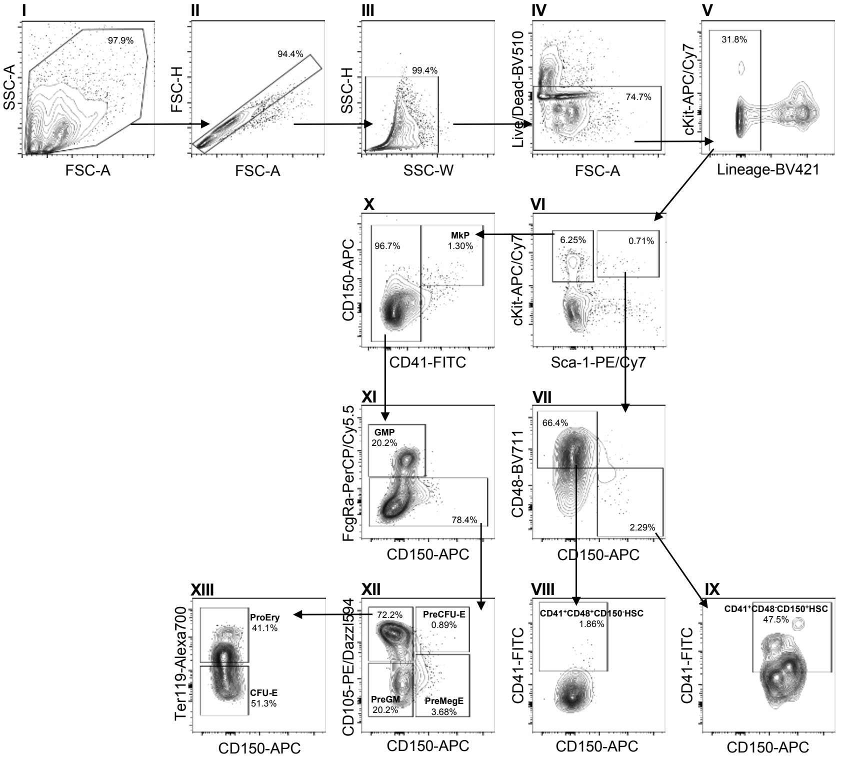


| Cell type | Immunophenotype |
|---|---|
| MP | Lin$^-$ Sca1$^+$ cKit$^-$ |
| MkP | Lin$^-$ Sca1$^+$ CD150$^+$ CD41$^+$ |
| GMP | Lin$^-$ Sca1$^+$ CD150$^-$ CD41$^-$ FcγRa$^+$ |
| PreGM | Lin$^-$ Sca1$^+$ CD150$^-$ CD41$^-$ FcγRa$^-$ CD105$^-$ |
| PreMegE | Lin$^-$ Sca1$^+$ CD150$^+$ CD41$^-$ FcγRa$^-$ CD105$^-$ |
| PreCFU-E | Lin$^-$ Sca1$^+$ CD150$^+$ CD41$^-$ FcγRa$^-$ CD105$^+$ |
| CFU-E | Lin$^-$ Sca1$^+$ CD150$^-$ CD41$^-$ FcγRa$^-$ CD105$^+$ Ter119$^-$ |
| PreEry | Lin$^-$ Sca1$^+$ CD150$^-$ CD41$^-$ FcγRa$^-$ CD105$^+$ Ter119$^+$ |
| LSK | Lin$^-$ Sca1$^+$ cKit$^+$ |
| CD41$^+$CD48-HSC | Lin$^-$ Sca1$^+$ cKit$^+$ CD48$^-$ CD150$^+$ CD41$^+$ |
| CD41$^+$ CD48$^+$HSC | Lin$^-$ Sca1$^+$ cKit$^+$ CD48$^+$ CD150$^{Nega}$ CD41$^+$ |

Lineage: B220, CD11b, CD3, Gr1, IL-7Ra
(CD11b was excluded for fetal liver samples)

**Yano_FigS3**

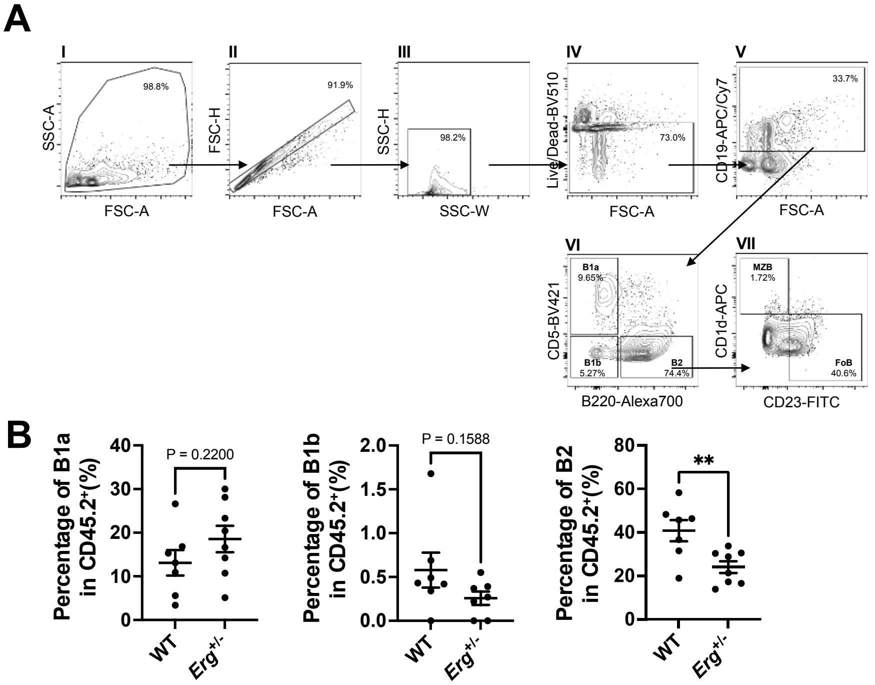
A
I
98.8%
SSC-A
FSC-A
II
91.9%
FSC-H
FSC-A
III
98.2%
SSC-H
SSC-W
IV
Live/Dead-BV510
73.0%
FSC-A
V
33.7%
CD19-APC/Cy7
FSC-A
VI
B1a
9.65%
B1b
5.27%
B2
74.4%
CD5-BV421
B220-Alexa700
VII
MZB
1.72%
FoB
40.6%
CD1d-APC
CD23-FITC
B
P = 0.2200
Percentage of B1a in CD45.2+(%)
WT
Erg+/-
P = 0.1588
Percentage of B1b in CD45.2+(%)
**
Percentage of B2 in CD45.2+(%)

Yano_FigS4

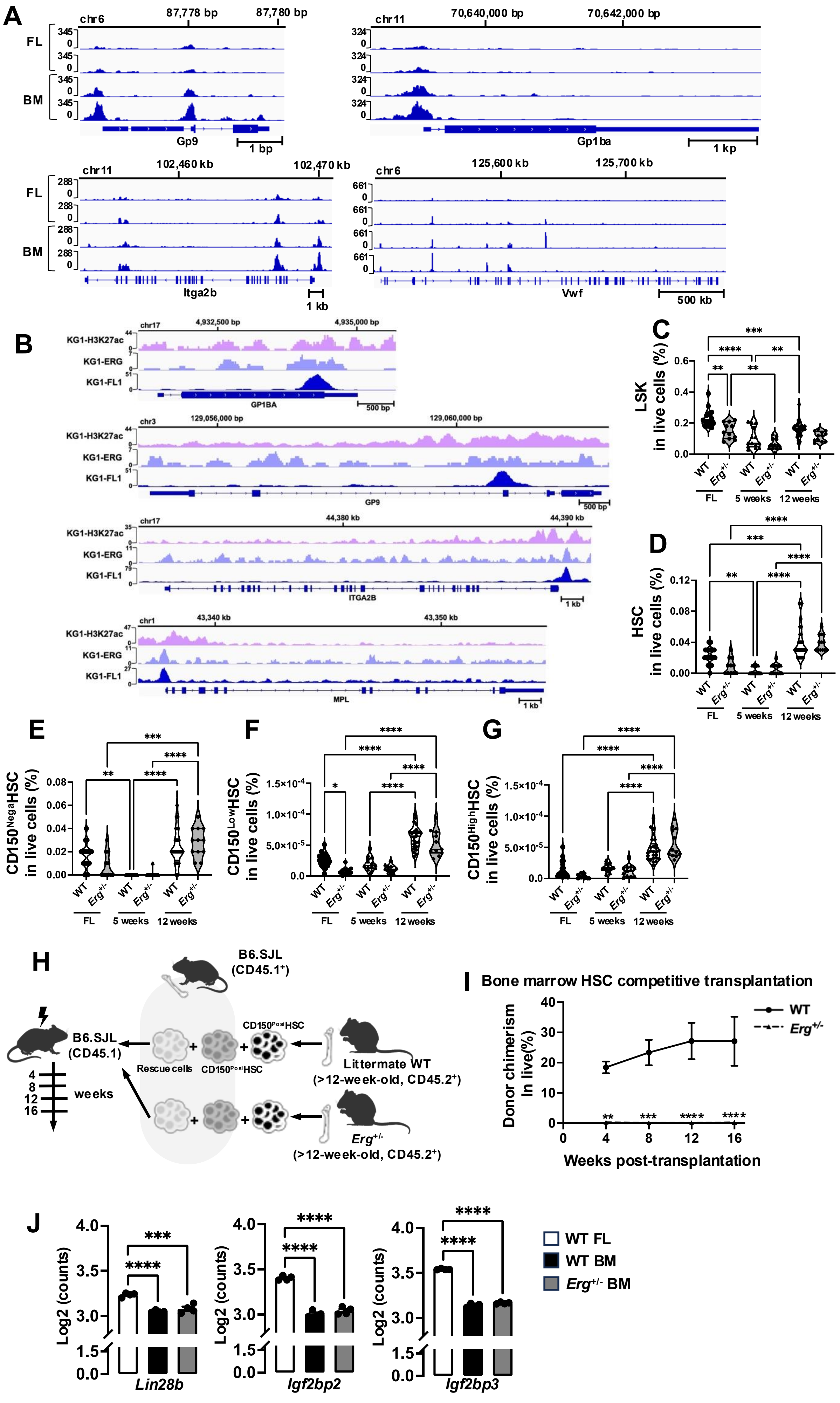
A
chr6
87,778 bp
87,780 bp
FL
BM
Gp9
1 bp
chr11
70,640,000 bp
70,642,000 bp
Gp1ba
1 kp
chr11
102,460 kb
102,470 kb
Itga2b
1 kb
chr6
125,600 kb
125,700 kb
Vwf
500 kb
B
KG1-H3K27ac
KG1-ERG
KG1-FL1
GP1BA
GP9
ITGA2B
MPL
C
LSK in live cells (%)
D
HSC in live cells (%)
E
CD150^Nega HSC in live cells (%)
F
CD150^Low HSC in live cells (%)
G
CD150^High HSC in live cells (%)
WT
Erg+/-
FL
5 weeks
12 weeks
H
B6.SJL (CD45.1+)
B6.SJL (CD45.1)
Rescue cells
CD150^Posi HSC
Littermate WT (>12-week-old, CD45.2+)
Erg+/- (>12-week-old, CD45.2+)
4
8
12
16
weeks
I
Bone marrow HSC competitive transplantation
Donor chimerism In live(%)
Weeks post-transplantation
WT
Erg+/-
J
Log2 (counts)
Lin28b
Igf2bp2
Igf2bp3
WT FL
WT BM
Erg+/- BM

### Table S1
### Hemavet data

| | | Litter control | *Erg*$^{+/-}$ | *P*-value |
|---|---|---|---|---|
| Total white blood cells | 5 wks | 8.19 K/μL ± 1.45 K/μL | 5.92 K/μL ± 1.16 K/μL | **$P$ = 0.0042 |
| | 12 wks | 8.90 K/μL ± 2.27 K/μL | 5.87 K/μL ± 1.55 K/μL | ****$P$ < 0.0001 |
| Neutrophils | 5 wks | 3.05 K/μL ± 0.76 K/μL | 1.11 K/μL ± 0.45 K/μL | *$P$ = 0.0155 |
| | 12 wks | 2.24 K/μL ± 0.76 K/μL | 1.59 K/μL ± 0.69 K/μL | **$P$ = 0.0051 |
| Lymphocytes | 5 wks | 4.01 K/μL ± 1.42 K/μL | 3.19 K/μL ± 1.04 K/μL | $P$ = 0.2743 |
| | 12 wks | 5.84 K/μL ± 1.55 K/μL | 3.68 K/μL ± 0.78 K/μL | **$P$ = 0.0051 |
| Monocytes | 5 wks | 0.60 K/μL ± 0.18 K/μL | 0.50 K/μL ± 0.14 K/μL | $P$ = 0.1948 |
| | 12 wks | 0.80 K/μL ± 0.28 K/μL | 0.58 K/μL ± 0.24 K/μL | *$P$ = 0.0194 |
| Red blood cells | 5 wks | 7.71 M/μL ± 0.43 M/μL | 8.23 M/μL ± 0.33 M/μL | **$P$ = 0.0042 |
| | 12 wks | 8.49 M/μL ± 0.90 M/μL | 8.89 M/μL ± 0.48 M/μL | $P$ = 0.1708 |
| Hemoglobin | 5 wks | 12.7 g/dL ± 0.43 g/dL | 13.2 g/dL ± 0.39 g/dL | *$P$ = 0.0316 |
| | 12 wks | 12.4 g/dL ± 0.42 g/dL | 12.3 g/dL ± 0.54 g/dL | $P$ = 0.7313 |
| Platelets | 5 wks | 919 K/μL ± 123 K/μL | 742 K/μL ± 93 K/μL | **$P$ = 0.0085 |
| | 12 wks | 928 K/μL ± 115 K/μL | 699 K/μL ± 113 K/μL | ****$P$ < 0.0001 |
